# Utilizing multi-objective decision support tools for protected area selection

**DOI:** 10.1101/2022.02.15.480531

**Authors:** Alke Voskamp, Susanne A. Fritz, Valerie Köcke, Matthias F. Biber, Timo Nogueira Brockmeyer, Bastian Bertzky, Matthew Forrest, Allie Goldstein, Scott Henderson, Thomas Hickler, Christian Hof, Thomas Kastner, Stefanie Lang, Peter Manning, Michael B. Mascia, Ian McFadden, Aidin Niamir, Monica Noon, Brian O’Donell, Mark Opel, Georg Schwede, Peyton West, Christof Schenck, Katrin Böhning-Gaese

## Abstract

The establishment and maintenance of protected areas (PAs) is viewed as a key action in delivering post-2020 biodiversity targets. PAs often need to meet multiple objectives, ranging from biodiversity protection to ecosystem service provision and climate change mitigation, but available land and conservation funding is limited. Therefore, optimizing resources by selecting the most beneficial PAs is vital. Here, we advocate for a flexible and transparent approach to selecting protected areas based on multiple objectives, and illustrate this with a decision support tool on a global scale. The tool allows weighting and prioritization of different conservation objectives according to user-specified preferences, as well as real-time comparison of the selected areas that result from such different priorities. We apply the tool across 1347 terrestrial PAs and highlight frequent trade-offs among different objectives, e.g., between species protection and ecosystem integrity. Outputs indicate that decision makers frequently face trade-offs among conflicting objectives. Nevertheless, we show that transparent decision-support tools can reveal synergies and trade-offs associated with PA selection, thereby helping to illuminate and resolve land-use conflicts embedded in divergent societal and political demands and values.

## Introduction

Halting biodiversity loss is one of the major global challenges faced by humanity in the 21^st^ century^1,2^. Human wellbeing, livelihoods, and economies all rely on biodiversity, and collaborative international efforts are needed to conserve it^1,3^. Protected areas (PAs) are a cornerstone of biodiversity conservation. Aichi Target 11 of the Convention on Biological Diversity called for an increase in PA coverage to 17% by 2020 for the terrestrial realm, with a focus on PAs that are of particular importance for biodiversity and ecosystem services, ecologically representative and well connected^4^; this goal has only partly been reached^5^. Further, Aichi target 11 is increasingly seen as inadequate to safeguard biodiversity^6–8^. The Kunming-Montreal Global Biodiversity Framework (GBF), which builds on the Aichi targets, has set out 23 action oriented global targets in line with an ambitious plan to implement broad action which should transform our societies’ relationship with biodiversity by 2030^9^. Action Target 3 of the GBF calls for at least 30 percent of the terrestrial area to be effectively conserved by PAs or “other effective area based conservation measures”^9^. This implies not only the transformation of large land areas into new PAs over the next decade, but also stresses an urgent need for careful allocation of the long-term conservation funding necessary to effectively protect biological resources: PAs must be both sustainably funded and effectively managed, yet only about 20% of all PAs are considered to meet these criteria^10^. Meanwhile, many PAs have experienced PA downgrading, downsizing or degazettement^11^ (PADDD) or are threatened by PADDD in the future^11,12^.

Both the allocation of sparse conservation funding for the strengthening of current PAs and the identification of additional sites to expand PA networks frequently require the application of prioritization approaches. A wealth of methods have been developed to inform conservation efforts, which vary widely in complexity. Some approaches evaluate individual sites based on their importance for the global persistence of biodiversity, e.g. the key biodiversity area (KBA) approach, applying different threshold-based criteria including the proportion of threatened or geographically restricted species covered^13^. In contrast, others rely on complex algorithms to optimize conservation networks towards specific conservation goals, e.g. by considering complementarity, connectivity, or cost efficiency^14–16^.

Priority areas for biodiversity conservation can be defined based on one or more individual conservation objectives, to identify areas of high conservation value under each or all given objectives. Initial approaches to identify such areas sought hotspots of various aspects of biodiversity such as species richness or endemism^17–20^. Other approaches highlight the protection of areas that will limit further impacts of global change on biodiversity, for example, by identifying remaining ecologically intact ecosystems^21^ or sites of high irrecoverable carbon storage^22,23^. Prioritization approaches that focus on more than one objective often combine different conservation goals like protecting biodiversity and maintaining ecosystem services. Here, we focus on those prioritization approaches that allow to identify individual sites of conservation importance rather than an optimized network of sites.

## The challenge: Aligning conservation priorities

Aligning different conservation objectives has become increasingly important. For instance, conservation strategies that address both ongoing climate warming and biodiversity loss are urgently needed^8,24^. Still, setting priorities based on multiple goals is not always straight forward. If there are trade-offs among conservation objectives, a very different set of sites might be optimal under each objective, and a simple compromise among these might not select the best set for the group of objectives as a whole. Relying on approaches tailored towards a single conservation objective, or the identification of one key element of the GBF targets, may lead to the omission of other critical elements of the GBF vision^25^.

To date, a vast amount of literature on setting global priorities for conservation is available (see Table S1 for an overview relevant to this study). The different approaches vary in the number of objectives that are considered, ranging from one to multiple, and the way the included variables are weighted, not all or with equal or uneven weights (Table 1). One of the earliest efforts to highlight global areas of importance for biodiversity protection are the global biodiversity hotspots identified by Meyers et al (2000)^26^. These were derived based the number of endemic species and habitat loss in the area. With the growing volume and availability of biodiversity data, more approaches to identify areas that are important for biodiversity protection have been introduced. Examples for individual or combined aspects of biodiversity that have been utilized for conservation priority maps are the global species richness patterns for terrestrial vertebrates or vascular plants as well as for various other taxonomic groups, but also biodiversity metrics such as species endemism, phylogenetic and functional diversity, or threat status have been used^27–31^. Similarly, increasing data availability and spatial resolution of those data has profited approaches that focus on prioritizing conservation sites based on the intactness of habitats and biomes or ecoregions^33^. Generally, priority maps for biodiversity protection can be derived based on a single metric for biodiversity or based on several combined metrics, as for example by combining the biodiversity value of an area with the level of threat, through human impacts like habitat degradation within the area^32,34^ (see Table S1 for more examples). Several efforts have also been made to align multiple conservation objectives, such as the protection of biodiversity, the preservation of ecosystem services and the preservation of areas important for climate mitigation. An example (Table 1) is the comparison of the spatial alignment of terrestrial biodiversity, carbon storage, and water quality regulation, and the identification of areas with the highest synergies among these objectives^35–37^. However, there is also evidence for trade-offs among conservation objectives, e.g. biodiversity hotspots do not always overlap with different ecosystem services^38^. In summary, a wealth of spatial prioritization maps for conservation efforts has been produced by all these different approaches, either to combine different biodiversity metrics to identify priority areas for biodiversity conservation or to align different conservation objectives to identify priority areas across these objectives. In fact, Cimatti et al (2012) subsequently combined 63 different global prioritization maps to derive one spatial prioritization map and identify scientific consensus regions among the different approaches^39^. Nevertheless, all of these selection approaches have one aspect in common: they result in a unique solution for one or a few specific and aligned objectives that selects a static geographic set of priorities (Table 1). Here, we advocate a more flexible approach that can handle multiple and conflicting objectives.

**Table 1:** A comparison of strengths and weaknesses of the approach advocated and implemented in this study vs. already existing approaches. The table summarizes a literature review, and gives a few selected examples from this. The review focused on studies that published global prioritization maps based on one or multiple conservation objectives and which identified individual sites of conservation importance rather than designed an optimized network of sites (see supplement and Table S1 for details and the considered studies).

| Approach | Methods (Tools) | Strength and weaknesses | Example studies | Objectives considered in the example studies |
| --- | --- | --- | --- | --- |
| *Single objective* | mapping | + Prioritization map based on one conservation objective<br>- Solution for one objective | Di Marco et al 2012<sup>43</sup>; Riggio et al 2020<sup>44</sup> | ecosystem integrity |
| *Multiple objectives* | mapping, stacked layers | + combined prioritization map across multiple objectives<br>- static solution, all objectives equally important | Jung et al 2021<sup>36</sup>; Dinerstein et al 2020<sup>8</sup> | biodiversity, ecosystem services, climate protection |
| *Multiple objectives + fixed weights* | mapping, stacked layers, consensus score | + combined prioritization map across multiple objectives<br>+ objectives (or variables within objectives) can be weighted individually<br>- static solution | Freudenberger et al 2013<sup>45</sup>; Girardello et al 2019<sup>46</sup> | biodiversity, ecosystem services, ecosystem integrity |
| *Multiple objectives + flexible weights* | mapping, stacked layers, weighted consensus score, individual ranking of sites | + combined prioritization map across multiple objectives<br>+ comparison of tradeoffs on the fly<br>+ flexible solution | ***This study*** | biodiversity, ecosystem integrity, climate protection, climatic stability, land-use stability, size |

The weaker the alignment is among different conservation objectives, the greater the influence of priority setting (i.e., favoring specific conservation objectives) on the outcome of site selection approaches. If trade-offs are prevalent, explicit values-based decision making is necessary. The relative priority of different conservation objectives varies among different societal groups, which differ in their demands and values^40^. Also, key local, national, and international actors – governments, corporations, non-governmental organizations (NGOs), scientists, and funders or sponsors – are likely to differ in their priorities^41^. Therefore, decisions as to which areas should be prioritized are often strongly values-based, with the values underlying final compromises rarely being made entirely explicit and transparent. Societal and political values are also likely to change over time, since the purpose of conservation itself has been transient over time, with priorities changing to some degree from one generation to the next^42^. All of this substantiates the need for a flexible but transparent approach to priority-setting, where different conservation objectives can be explicitly considered and weighed against each other, to facilitate deliberative societal and political decision making.

## Towards a solution: flexible and transparent site selection

The allocation of conservation funding is one example where the use of a flexible and transparent prioritization approach can be advantageous since the decision process is likely to involve multiple stakeholders, each of which may have multiple objectives. Use of a decision support tool can support the identification of conservation synergies and trade-offs, facilitate deliberation and dialog among stakeholders, and enable evidence-informed, values-based collaborative decision-making. Here, we illustrate these ideas using a site selection tool that we developed for this task. We apply a transparent site selection approach that allows users to identify investment priorities among existing PAs based on various self-specified conservation objectives. In contrast to other approaches, conservation objectives in our approach are explicitly weighted by the users and the results can be immediately assessed, aiding discussions during a transparent values-based decision-making process. We implemented the approach for the terrestrial realm, exclusively using biogeographic information that is publicly available at a global scale. We aimed to identify areas with the highest potential for a range of biodiversity and climate protection goals, but excluded any information on political and economic dimensions from the site selection algorithm; although these considerations are crucial for conservation and should be evaluated equally transparently, we believe that they should be evaluated separately from biogeographic information as an additional step in the decision-making process.

We defined six different conservation objectives (Fig. 1), which represent a broad agreement on priorities for safeguarding biodiversity, climate protection (in the sense of mitigating ongoing climate change), and the present and projected future status of individual sites (identified in an initial stakeholder dialog, see also case study details below). These objectives were: 1) high current biodiversity, focusing on high biodiversity values, 2) high current ecosystem integrity, which focuses on areas that have experienced relatively few anthropogenic impacts, 3) high climate protection, which selects for sites that have large, irreplaceable carbon stocks, 4) large size, which prioritizes bigger sites, 5) high land-use stability, which focuses on the future likelihood of land-use change in the immediate surroundings of sites, and 6) high climatic stability, which highlights sites in which climate change is projected to have low impacts on current biodiversity.

**Fig. 1:**
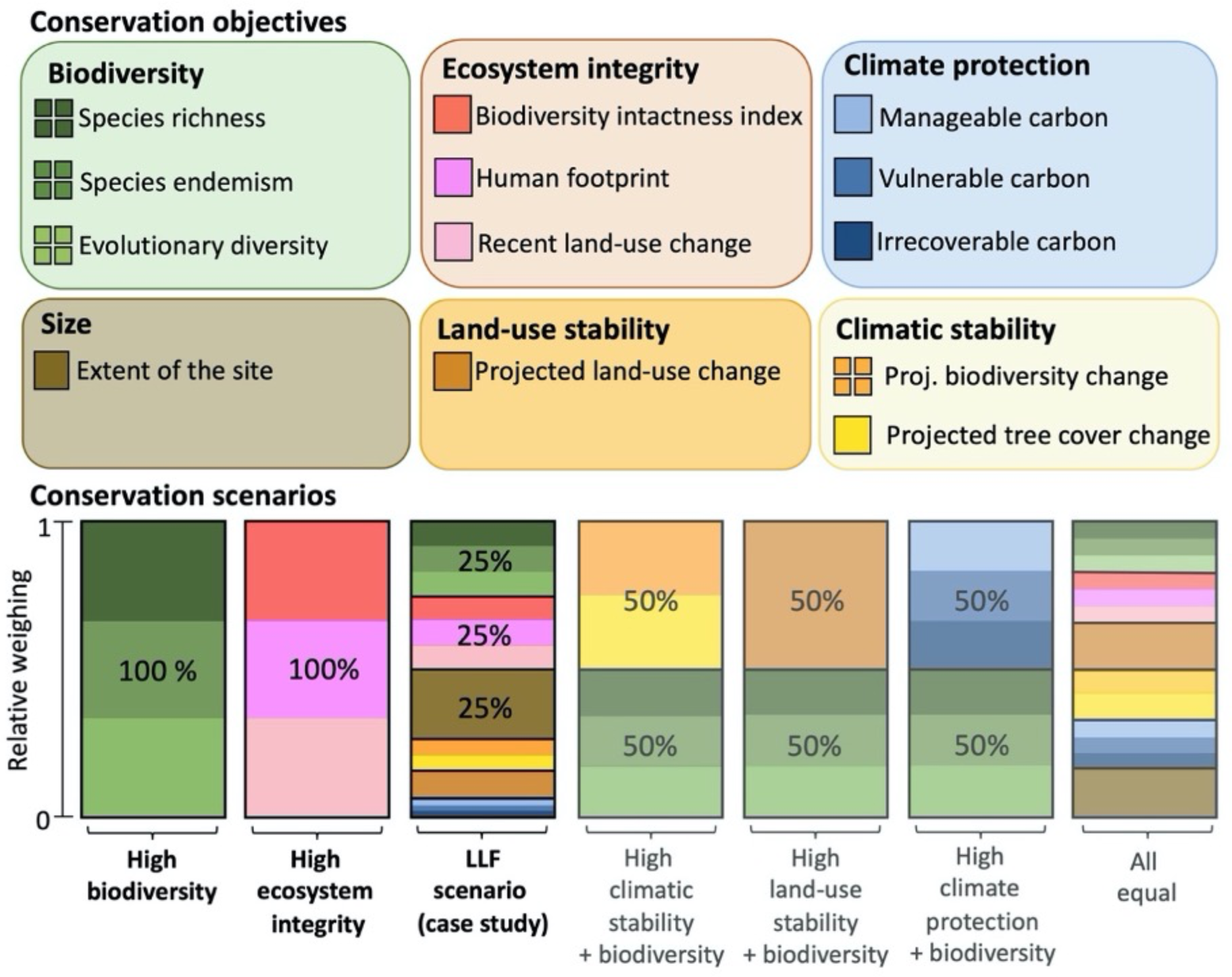
The six conservation objectives defined to set priorities for the site selection, the indicators considered for each objective (note that Biodiversity and Climatic stability (of biodiversity) include indicators for four different vertebrate taxa), and examples for conservation scenarios based on these objectives. By applying a weighting approach, user-specified objectives can be combined into different conservation scenarios, which are therefore customized for specific conservation goals. The High biodiversity, High ecosystem integrity and Legacy Landscapes Fund (LLF) scenarios are used in the case study.

We collated a broad set of conservation indicators that reflect these six conservation objectives (Fig. 1). The biodiversity objective considered as indicators the total terrestrial species richness of four vertebrate taxa (birds, mammals, amphibians and reptiles) as well as species endemism and evolutionary diversity^47^ for each taxon, to capture the amount of biodiversity as well as its irreplaceability. The ecosystem integrity objective considered biodiversity intactness, recent land-use change, and the human footprint within the site. The climate protection objective considered the average amount of carbon per hectare that is stored in the vegetation and soil (up to 1 meter below ground) of the site and its vulnerability to typical land conversion. The size objective covers the extent of the site in km^2^. The land-use stability objective considered the projected change in land-use in a buffer zone around the site. The climatic stability objective considered the biodiversity change based on the projected future compositional change (turnover)^48^ of the four vertebrate taxa and the projected change in tree cover within the site.

These conservation objectives and the underlying indicators were carefully selected reflecting the demands towards the PA network based on the post-2020 GBF, as well as the current state of the literature addressing both the biodiversity and climate crises. The biodiversity objective combines information on the number, diversity and rarity of species across several higher taxa within the area, to include different aspects of biodiversity^47,49–52^. Highlighting those sites that are of particular importance for biodiversity is in line with the first part of Action Target 3 of the post-2020 GBF^9^. The ecosystem integrity objective uses information on recent impacts on the site and the intactness of the local ecological communities, highlighting those sites that contain ecosystems that are still largely intact. This objective was included because remaining intact ecosystems are often not directly addressed by conservation efforts or international policy frameworks^21,53^, but provide various key functions, such as acting as critical carbon sinks, stabilizing hydrological cycles, or providing crucial refuge for imperiled species, intact mega-faunal assemblages, or wide-ranging or migratory species^21,54–59^. The size objective is somewhat related to the ecosystem integrity objective, under the assumption that larger areas have a higher potential to support populations of target species and to maintain functioning ecosystems in the long term^60,61^. The climate protection objective is related to Action Target 8 of the post-2020 GBF, which aims to minimize the impacts of climate change on biodiversity.

The final two objectives were included to assess sites not only based on their current importance for biodiversity, ecosystem functioning, and climate protection, but also based on the most major future threats towards biodiversity, i.e. projected future climate and land-use change. The five direct drivers of biodiversity loss with the largest impact, according to the 2019 Global Assessment Report by IPBES, are changes in land and sea use; direct exploitation of organisms; climate change; pollution; and invasion of alien species^1^. The climatic and land-use stability objectives provide an indication of potential future changes within the site based on climate change responses (geographic range shifts) of the local flora and fauna within the region and give an indication of which sites might be under increasing pressure of land-use change in the region.

A key aspect in developing a transparent site selection approach was to make results of different values-based objective weighting immediately accessible to a broader audience, including decision makers. We therefore developed an open-source spatial decision support tool to facilitate the priority-based area selection process. The tool generates a ranking of sites globally as well as for each biogeographic realm, based on the six conservation objectives which are weighted individually by the user. Using sliders to allocate weights to the six conservation objectives, users can design their own conservation scenarios on the fly (examples see Fig. 1), and directly visualize the resulting ranking. The tool allows a comparison of a far wider range of different conservation scenarios than the examples we give here, to evaluate synergies and trade-offs among these, and select sites for a more detailed investigation. The current version is publicly available (https://ll-evaluation-support-tool.shinyapps.io/legacy_landscapes_dst/) and restricted to the case study dataset, objectives and indicators presented in the paper, but the flexible approach we use can be implemented easily to other datasets, objectives, and goals.

## Illustration of the selection approach: The Legacy Landscapes Fund as a case study

The Legacy Landscapes Fund (LLF) is a recently established foundation that provides long-term funding for protected areas^62^; it is useful in this context because it uses our six conservation objectives, operates on a global level, and mostly focuses on existing sites. This allowed us to run a case study across a significant set of PAs and other sites of interest across the globe, in order to demonstrate how the newly developed decision support tool facilitates the flexible evaluation of potential priority sites for conservation and to explore the potential and limitations of this approach. We assessed synergies and trade-offs among areas according to the different objectives at a global scale, as well as within biogeographic realms. Finally, we aimed to investigate how priority setting by different societal actors affects site selection by combining the multiple conservation objectives into broader conservation scenarios that weigh each objective according to user-specified priorities.

The case study dataset for the analysis contained 1347 sites globally. These sites included formally protected areas of IUCN category I or II, listed Natural World Heritage Sites (WHS) and registered Key Biodiversity Area (KBA) (see experimental procedures and supplementary material for details on dataset and methods)^63,64^. A principal component analysis (PCA) applied to this dataset globally (Fig. 2) and at the level of biogeographic realms (Fig. 3) showed that the indicators belonging to each conservation objective tended to be closely aligned both at the global and the realm level, with the only exception being the two climatic stability indicators across the Australian realm. For example, within the biodiversity objective, species richness (SR), species endemism (SE) and evolutionary diversity (ED) were closely aligned at the global scale, as well as at the biogeographic realm level, though the alignment between SR and the other two indicators was slightly less tight in the tropical realms (Fig. 3).

**Fig. 2:**
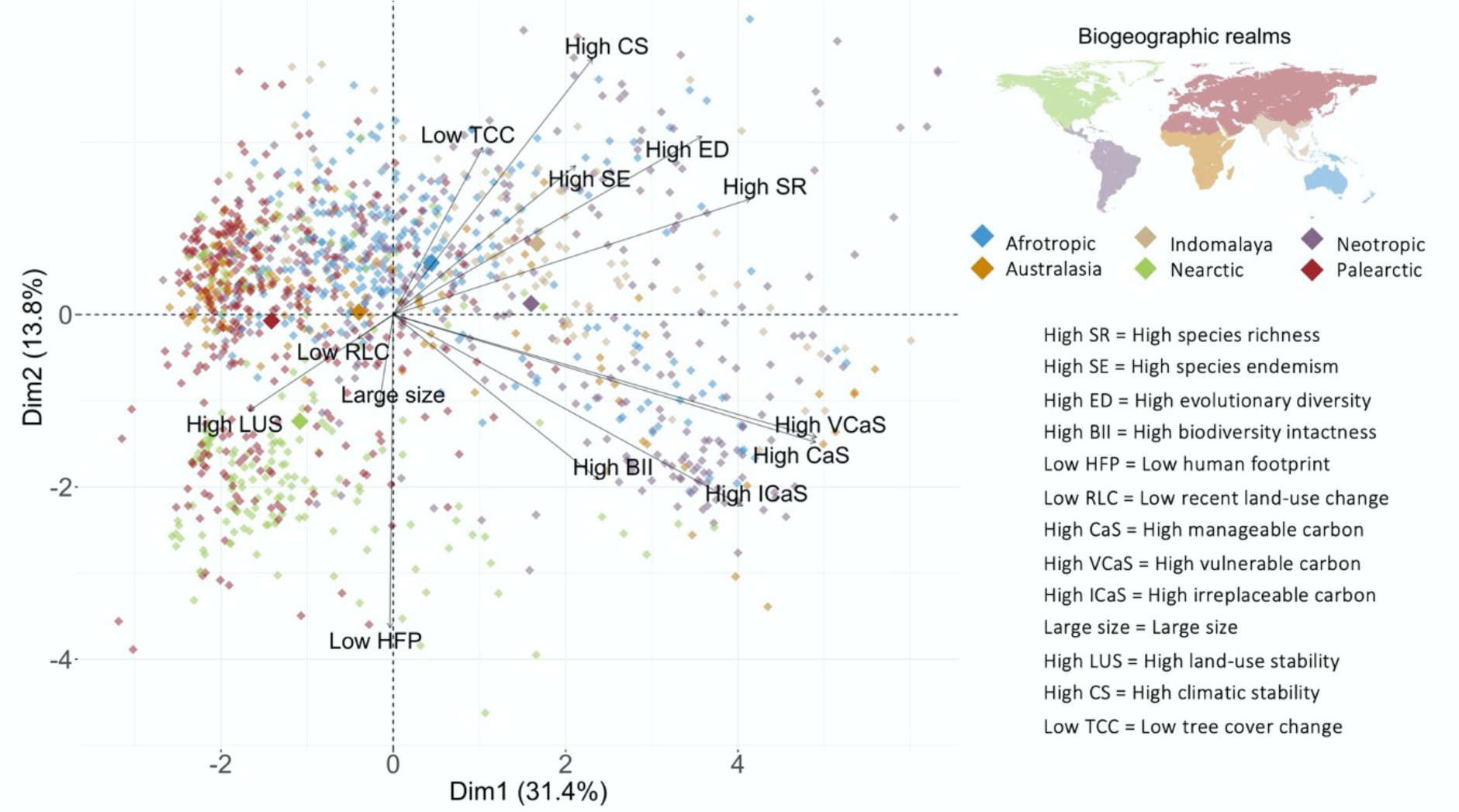
Trade-offs and synergies between the conservation indicators of individual sites. Shown are the first and second dimensions of a principal component analysis (PCA) that was performed across 1347 sites and their variation in 13 indicator variables aggregated into six conservation objectives (order of indicator variables in the legend aligns with Fig. 1 and 3, see these for matching variables to objectives). The first and second PCA dimensions together explain 45.3% of the variation in the data. Each dot represents one site. The arrows represent the indicators and the arrow length indicates the loading of each indicator onto the PCA dimensions (i.e. their correlation with each principal component). Opposite loadings indicate trade-offs between the variables (i.e., a site that has a high value in one of these variables, has a low value in the other variable and vice versa). The individual sites (points) are colored by the biogeographic realm in which they are located^65^.

**Fig. 3:**
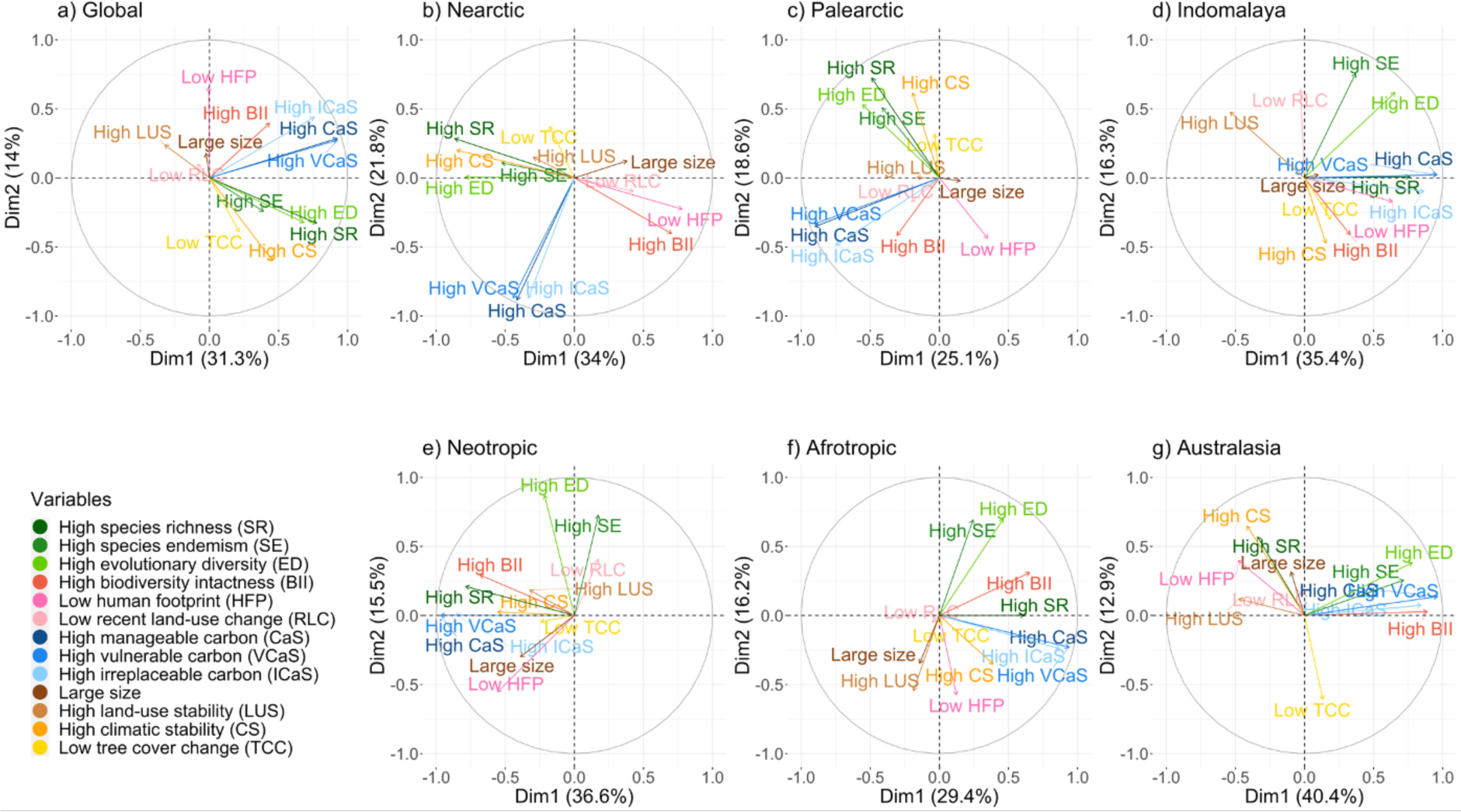
Trade-offs and synergies between the conservation indicators of individual sites at the global and realm levels. Shown are the first two axes of the principal component analysis (PCA) for all 1347 sites included in the Legacy Landscapes case study globally and for each individual realm. These analyses reveal trade-offs between the conservation objectives, indicated by variables mapping onto opposing ends of a principal component axis. Variable colours indicate conservation objectives as in Fig. 1: biodiversity (shades of green), ecosystem integrity (shades of red and pink), climate protection (shades of blue), size (dark brown), land-use stability (light brown) and climatic stability (orange and yellow). PCA plots show the respective first two axes identified and the percentage of variation explained by each of the axes.

Looking at the trade-offs and synergies among the objectives, we found that at the global scale the first and second PCA axes explained 31.4 and 14.2 percent of the variation in the data respectively. These axes showed relatively clear trade-offs and synergies among the six different conservation objectives (Fig. 3). The strongest global trade-off was found between current biodiversity and future land-use stability (Pearson’s correlation coefficient r (n=1346) = −.30, p<0.01). These two objectives are negatively correlated, as increasing land-use pressure is often projected to occur around sites with exceptionally high current biodiversity (e.g. deforestation of tropical forests for agriculture). The strongest global synergies were found between current biodiversity and future climatic stability (r (n=1346) = .41, p<0.01) and current biodiversity and high climate protection potential based on the amount of manageable carbon stored in the site (r (n=1346) = .58, p<0.01). This suggests that sites with exceptionally high biodiversity often coincide with areas of lower projected impacts of climate change on vertebrate communities and tree cover and with a high potential for climate protection through carbon storage. The identified global synergies and trade-offs between the different objectives were only partially consistent within realms, with patterns very similar to the global analysis for the Afrotropical realm but notably different alignments in the Palearctic and Nearctic.

Finally, to investigate how priority setting by different societal groups can affect site selection, we compared the outcome of area selection under three different conservation scenarios. We used two extreme and one combined scenario, to explore a broad range of values (Fig. 1). The first scenario was a biodiversity scenario (biodiversity objective weighted by 100% and the other five objectives by 0%). The second was an ecosystem integrity scenario (ecosystem integrity 100%, all others 0%). The third scenario was a stakeholder-driven scenario that resulted from joint discussion during an expert workshop (LLF scenario; Fig. 1). At this two-day online workshop, which was attended by 35 experts with a strong conservation background, we introduced the site selection approach, further developed the indicators and objectives, and voted on the LLF scenario (see supplementary materials for more detail). This scenario reflects the main selection criteria for potential LLF sites (high biodiversity, ecosystem integrity and size) but considers also the other objectives weighted according to lower priorities (biodiversity, ecosystem integrity and size weighted with 25% each, climatic stability and land-use stability with 10% each, and climate protection with 5%).

Despite synergy between some objectives, we found that when comparing the top five sites selected for each of the three conservation scenarios, within each biogeographic realm, there is little congruence among these scenarios (Fig. 4). This implies that selecting sites based on their biodiversity will in most cases result in the protection of different sites compared to a selection based on high ecosystem integrity, or the LLF scenario. Australasia has the highest overlap of top sites for the three different scenarios, with four sites being in the top five for both the biodiversity and the LLF scenario. The Nearctic, Neotropic and Afrotropic realms have the least overlap among the top sites for the investigated scenarios with only one shared site in the top five of all scenarios.

**Fig. 4:**
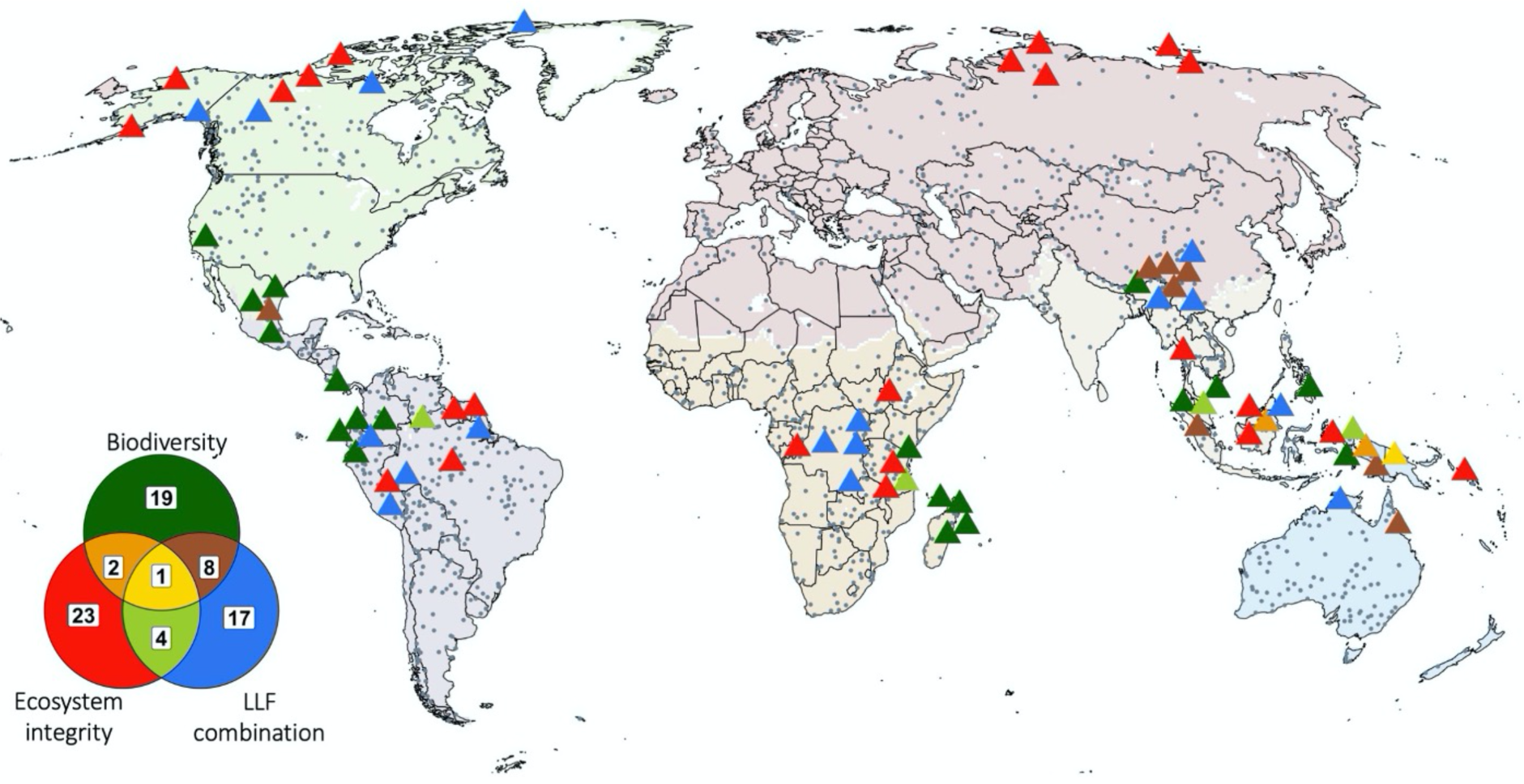
Spatial distribution of sites highlighting the top 5 priority sites for each of the 3 example conservation scenarios: prioritizing biodiversity (dark green), prioritizing ecosystem integrity (red) and the LLF scenario (Legacy Landscapes Fund, prioritizing a combination of all objectives that stresses high biodiversity, high ecosystem integrity and large size; blue). The top 5 sites for all three scenarios (triangles) are shown per biogeographic realm (i.e., 30 top sites per conservation scenario in total). The colors correspond to the three different conservation scenarios and their overlap (if a site is in the top five for more than one objective), as shown in the Venn diagram. Only 14 of the top sites were selected under two scenarios (light green, brown and orange) and 1 site was selected under all 3 scenarios (yellow). Grey points indicate sites included in the analysis but not selected under the top 5. Top sites in close geographic proximity are spaced out for visualization and deviate from their exact spatial position. Map colors indicate the different biogeographic realms.

## Discussion

Our case study demonstrates that the selection of ‘best’ sites for nature conservation depends largely on the relative weighting of different conservation priorities and is therefore heavily influenced by decision-maker values. This is supported by the clear trade-offs among the six conservation objectives at the realm and global scale (Fig. 2, 3), as well as the limited congruence among the top sites selected under the three different conservation scenarios (Fig. 4). These results illustrate the opportunities and challenges faced by decision makers when selecting priority areas for nature conservation. Furthermore, they demonstrate the need for a global approach to nature conservation that involves multiple stakeholder groups and perspectives and a transparent decision-making process. Here, we introduce an approach to select priority areas for biodiversity conservation at the global scale that separates 1. global biogeographic information on biodiversity, ecosystem services, etc., from 2. a value-based prioritization of different conservation objectives in the decision-making process. This allows the trade-offs between conservation objectives to be understood and acknowledged explicitly and quantitatively. It thereby enables a first transparent evaluation of sites that reflects the varying priorities among different societal or conservation actors. Furthermore, the approach allows to optimize site selection towards more than one objective, which can significantly increase the efficiency of a PA network^66^. Additionally, the transient nature of conservation goals or new drivers of biodiversity loss, such as climate change, might result in the need to adjust prioritization in the future. Both arguments highlight the advantages of a flexible site selection approach over the static selection of hotspots based on a small number of fixed objectives and indicators.

Our approach goes beyond existing studies that explore the spatial agreement of conservation objectives and present optimized solutions through aligning several objectives, by allowing the user to change the prioritization on the fly (Table 1). Instead of presenting a static conservation priority map, we present a dynamic result that ranks potential sites for protection based on user preferences. This approach puts the focus on the decision making process and allows the exploration of tradeoffs and synergies among different options. Rather than providing another method to set conservation priorities, our approach is complementary to the various approaches we found in the literature (Table 1 and S1). It could for example be used to explore the differences, synergies and tradeoffs between any of the existing global prioritization maps, across protected areas.

### Applying the tool to a specific conservation problem

For the Legacy Landscapes Fund, the three conservation objectives of size, biodiversity and ecosystem integrity are of high priority^67^. Applying the decision support tool to the assembled dataset revealed a trade-off between high biodiversity and high ecosystem integrity, clearly demonstrated in the comparison between the three conservation scenarios: high biodiversity, high ecosystem integrity and the LLF scenario, which considers multiple conservation objectives. For the actual area selection to be financed by the LLF, the decision support tool enabled an initial screening of potential sites globally, to evaluate the performance of individual sites under the desired conservation objectives and to compare different weightings before proceeding with the selection of the pilot sites. Here, the decision support tool was used in an integrative decision-making process which transparently separated biogeographical site screening from other criteria like stakeholder consent, political commitment, and experience of the implementing NGO (also see below).

### Applying the approach beyond the case study

Our approach and the newly developed tool can be easily extended to include a broader range of biogeographic datasets, additional conservation objectives, or additional sites into the analysis, making the tool widely applicable to a variety of site selection tasks. Though the current set-up of the tool already contains six objectives representing several broad conservation goals (i.e. safeguarding biodiversity or mitigating climate change), these are still to some extent geared towards the case study. To broaden the scope of the tool through additional objectives and opposing the focus on intact ecosystems used in our case study, priority setting could highlight areas that harbor a high amount of threatened biodiversity^68^, e.g. by including an additional objective based on the threat status of all occurring species (i.e. as provided in the IUCN Red List) in a site^49,69,70^. Another obvious and easy possibility to expand the current set-up of the tool would be to allow further subsetting of the included sites. Currently the tool allows for an initial screening of sites at the level of biogeographic realms or at the global scale. Information such as the extent of a biogeographic realm or ecoregion that is already protected would need to be considered separately. Adjusting the tool to rank sites not only at the realm level but also at finer scales, as for example at the ecoregion level, would allow users to prioritize sites in finer-scale underrepresented categories.

Action Target 8 of the post 2020 GBF also calls for a well-connected PA network^9^. Connectivity is highly species-specific and landscape-dependent, and thus requires local and long-term studies on individual species^71,72^. Assessments on a scale like the decision support tool shown here cannot yet assess connectivity at that level. Still, previous efforts have estimated the connectivity of global PA networks at a coarser scale, for example based on different levels of home range size in mammals^73^ or even by modeling the movement of large animals throughout the landscape between protected areas^74^. A first step to integrate connectivity into the decision support tool could be to use a distance matrix of sites from surrounding existing PAs. This could give a first rough indication of how well a site is embedded into the PA network and allow prioritization of connected sites over very isolated sites.

As currently designed, the tool is meant to allow the comparison of sites and different conservation objectives based on biogeographic variables, which are available at a global scale. This necessitates the use of relatively coarse-grained datasets (resolution here is mostly dependent on the biodiversity data). The tool allows an initial screening of a large number of potential sites globally (or regionally) and can be extremely useful in creating prioritizations of PAs based on different objectives and indicators that can be applied flexibly. This tool, however, is only useful as a first step that allows a range of options to be explored, as part of a much broader decision-making process. This decision-making process should include on-site assessments of additional parameters at a higher resolution (e.g. more detailed biological data acquired through surveys and observations) as well as non-biological characteristics. These socio-economic factors could include, for example, the political legitimacy of the initiative, the involvement of local communities, and the presence of a supportive NGO. In case of pilot site selection for the LLF, these factors were considered in the next step that followed the use of the site evaluation tool. Further, the decision support tool was designed to facilitate value-based discussions by enabling on-the-fly comparison of sites based on different biogeographic attributes. The tool does not facilitate the optimization of site networks (i.e. assess different combinations of sites based on representativeness or cost efficiency).

### Applying the decision support tool within the post-2020 Global Biodiversity Framework

The ambition of the Aichi Biodiversity Targets has been increasingly criticized as being too modest to safeguard biodiversity in perpetuity^6,7^. Accordingly, the post-2020 GBF of the Convention on Biological Diversity calls for ‘at least 30 per cent of terrestrial, inland water and of coastal and marine areas, especially areas of particular importance for biodiversity and ecosystem functions and services, to be effectively conserved’^9^. Thus it becomes increasingly important to identify new sites for conservation – and new ways of conserving – outside of the already delineated areas both on land and in the oceans^8,75^. The presented decision support tool could be extended to aid these efforts, either by adapting it to identify new sites or by expanding the case-study dataset. A first possible extension would be the inclusion of the not yet formerly recognized Indigenous and Community Conservation Areas (ICCAs) and of Other Effective Area-based Conservation Measures (OECMs) which are increasingly being recognized as effective and potentially more inclusive conservation tools^76^.

Going beyond global priority-setting, the post-2020 GBF aims to facilitate implementation primarily through activities at the national level. Furthermore, unlike in the LLF case study, a vast amount of conservation funding is not available at the global scale but rather at the national or regional level. Our approach could be used at the national or sub-national level to help prioritize conservation decisions through facilitating transparent value-based discussion and support implementation of the post-2020 GBF at this scale^77^. Applying the tool at the national or regional scale would open the possibility to add more finely resolved datasets to the conservation objectives that are not available at the global scale (for example, species abundances or more specific land-use projections) and thus tailor the decision support tool to specific conservation actions.

An example of a relevant adjustment that may be possible at national scales could be the adjustment of the intended timeframe, as the decision support tool with its inclusion of future projections (climatic and land-use stability) as well as the focus on intact ecosystems is currently geared towards longer time horizons. Highlighting sites where there is an urgent need to act (e.g. within a couple of years because of high conservation value in combination with high current pressure) would require the use of very different datasets with a much higher resolution. Working at regional or national scales would allow the inclusion of data sets on recent changes within a site that are not available or very heterogeneous at the global scale (e.g. population trends, recent deforestation rates, or the level of exploitation of natural resources).

In conclusion, the proposed approach facilitates a transparent initial screening of potential priority sites that allows the trade-offs between conservation objectives to be understood and acknowledged explicitly and quantitatively. It promotes the inclusion of multiple stakeholder positions, views and preferences, and facilitates discourse and decision-making whilst working towards the overarching conservation goals.

## Experimental procedures

### Lead contact

### Materials availability

This study did not generate unique new materials.

### Data and code availability

All codes needed to replicate the presented analysis are available from GitHub (https://github.com/Legacy-Landscapes/LL_analysis). The decision support tool is accessible via: https://ll-evaluation-support-tool.shinyapps.io/legacy_landscapes_dst/). All codes for the decision support tool are available under https://github.com/Legacy-Landscapes/LL_Decision_Tool.

### Conservation objectives data

The six defined conservation objectives are each based on several underlying data sets, with more detail on variable calculations and score assignations for each objective given in the supplement. The datasets behind the biodiversity objective are the global range-map polygons for all terrestrial birds, mammals, amphibians and reptiles as provided by BirdLife International, IUCN and GARD^49–51^, as well as the phylogenetic supertree for all four terrestrial vertebrate taxa from Hedges *et al*. 2015^78^. From these datasets we derived species richness, species endemism (calculated as corrected range size rarity^52^) and phylogenetic endemism^47^ values per site included for all four vertebrate taxa. The datasets underlying the ecosystem integrity objective are the biodiversity intactness index^79^, the human footprint compiled by Venter et al 2016^80^ and the recent land-use change 1992-2018 derived from the ESA CCI Land Cover by Niamir et al 2020^81^. The climate protection objective consists of three different indicators, the amount of manageable carbon stored in the site, the amount of vulnerable carbon and the amount of irrecoverable carbon^22,23^. The size objective uses the size of each site derived in QGIS^82^. All future stability variables were derived by comparing the timespan between 1995 (average climate projections 1980 – 2009) and 2050 (average climate projections 2035 – 2064). The climatic stability objective consists of two main underlying indicators, the climatic stability of biodiversity and the projected tree cover change. The climatic stability was calculated based on modelled changes in species community compositions that resulted from projected range shifts under climate change for all four taxa^83^. The projected change in tree cover is based on the LPJ-GUESS process-based dynamic vegetation-terrestrial ecosystem model^84^. Finally, the land-use stability objective consists of projected changes in five different land-use types (rainfed crop, irrigated crop, pastures, as well as rainfed and irrigated bioenergy crops), based on the MAgPIE and REMIND-MAgPIE model^85–87^ and using the assumptions of population growth and economic development as described in Frieler et al 2017^88^. These projections are based on the same climate projections as the climatic stability variables.

The six conservation objectives were developed in a discussion process among the broader conservation community. We introduced our approach at a two-day webinar which was attended by 35 experts with a strong conservation background. These included 1) conservation scientists, 2) international conservation NGOs, 3) the financial sector, and 4) policy sectors, in particular the German Federal Ministry for Economic Cooperation and Development (BMZ). These experts provided feedback on the objectives and indicators through a questionnaire (see supplementary material). They were asked to: 1) report any missing objectives, 2) report any missing indicators that should be included in the objectives and 3) rank the suggested objectives by their personal preferences. To translate personal preferences into site selection, the resulting ranks for each individual indicator were scaled from zero to one. Each objective consists of several underlying indicators (datasets), so by taking the mean across all indicators per objective these were weighted equally.

### The case study dataset and analysis

To assess synergies and trade-offs among the conservation objectives, we used the LLF as a case study to assemble a global dataset of sites. The LLF is a recently established foundation that provides long-term funding of one million U.S. dollars per “legacy landscape” per year. Funding stems from public and private sources. It aims to protect areas of outstanding biodiversity over initially 15 years – but with a vision to ensure funding in perpetuity^67^. The LLF is based on a strategic global site-selection approach and the strong long-term commitment of local NGOs, protected area authorities and local communities ‘on the ground’^62^. The initial requirements for sites to be considered by the LLF are outstanding biodiversity, a minimum size of 2,000km^2^ and a protection status as IUCN protected area category I or II for at least 1,000 km^2^. Based loosely on these guidelines, we assembled a dataset and extracted site-specific values for each objective (Fig. 1) (see supplementary material for a detailed account how the site dataset was assembled).

We then investigated global synergies and trade-offs among the final set of conservation objectives using a principal component analysis (PCA) across sites. To further explore if synergies and trade-offs between the objectives were different in biogeographic regions of the world, we repeated the PCA separately for each of the six terrestrial biogeographic realms^65^. Additional analyses are described in the supplement.

### The decision support tool

To make the analysis accessible to the broader conservation community and to enable a rapid comparison of sites based on the user-specified prioritization of the different conservation objectives, we designed an interactive spatial decision support tool in which weightings can be modified (see supplementary material for detailed content of the app interface). The user interface for the tool was developed using R Shiny version 1.5.0^89^.

## Acknowledgements

We thank all participants of the expert workshop for valuable discussions and input to the development of the decision support tool, and the FZS staff in the project areas who tested the tool and helped to evaluate its use for conservation. We gratefully acknowledge the use of the Goethe-HLR HPC at the Centre for Scientific Computing at Goethe University Frankfurt for some of the computationally heavy aspects of this work. We also thank BirdLife International for making the KBA data available as well as the Inter-Sectoral Impact Model Intercomparison Project (ISIMIP) and ISIpedia - the open climate-impacts encyclopedia for their support and data availability.

## Author contributions

Conceptualization: AV, SAF, VK, CS and KBG; Methodology: AV, SAF, KBG; Feedback on Methodology: all authors, Software: AV, TNB, MFB; Writing – Original: AV, SAF, VK and KBG; Writing – Review and Editing: all authors, Supervision: SAF and KBG.

