## Supplementary material for "Utilizing multi-objective decision support tools for protected area selection"

### Content

**1. SCREENED LITERATURE ON GLOBAL PRIORITIZATION APPROACHES ... 3**

**2. SUPPLEMENTARY METHODS AND MATERIALS ... 10**

2.1 THE PROTECTED AREA DATASET ... 10
2.2 THE CONSERVATION OBJECTIVES ... 11
*2.2.1 Biodiversity ... 11*
*2.2.2 Ecosystem intactness ... 13*
*2.2.3 Climatic stability ... 14*
*2.2.4 Land-use stability ... 16*
*2.2.5 Climate protection ... 16*
*2.2.6 Large size ... 18*
2.3 SCALING AND WEIGHTING THE INDICATORS FOR THE SITE EVALUATION ... 23
2.4 THE PRINCIPAL COMPONENT ANALYSIS (PCA) ... 23
2.5 SENSITIVITY OF SITE RANKINGS ... 24
2.6 THE WEBINAR ... 28

**3. CAVEATS ... 32**

**4. THE DECISION SUPPORT TOOL ... 33**

4.1 USER MANUAL DECISION SUPPORT TOOL ... 39
*4.1.1 Details on the conservation objectives ... 39*
*4.1.2 How to use the conservation decision support tool ... 44*

**5. REFERENCES ... 48**

### 1. Screened literature on global prioritization approaches

At the start of the project, we screened the available literature on global prioritization approaches, to identify a suitable tool that would allow to explore and compare the trade-offs between different conservation objectives flexibly. Although there are numerous prioritization approaches available based on various conservation objectives, none of them was applicable for the task at hand. The majority of approaches presented static maps on global priority areas for conservation based on one or more objectives, and only very few approaches considered weighing the included objectives (or variables included within the objectives) to obtain a consensus map across the different objectives. Table S1 provides a list of the global studies that we selected as highly relevant to our approach (i.e. not those with a focus on prioritizing a network of sites or identifying sites of high complementarity to an existing site network, but rather studies that resulted in priority maps across assemblages, sites, or some other spatial unit).

**Table S1:** Studies that present global prioritization maps based on one or multiple conservation objectives. The column objectives considered shows if the study is focused on a 'single' objective, which could be based on one or more variables (e.g. biodiversity measured based on several indicators like species richness, number of threatened species, etc.); on 'multiple' objectives which could be based on several variables (e.g. biodiversity, measured based on several indicators like species richness and number of threatened species, as well as ecosystem integrity, measured based on several indicators such as human footprint and intactness of species assemblages); or on 'multiple weighted' objectives which could be based on several variables and where the objectives (or the variables within the objectives) were not equally weighted.

| # | Authors | Year | Title | # of objectives considered | Variables |
|---|---|---|---|---|---|
| 1 | Albuquerque et al | 2015 | Global patterns and environmental correlates of high-priority conservation areas for vertebrates | single | vertebrate richness complementarity |
| 2 | Allan et al | 2022 | The minimum land area requiring conservation attention to safeguard biodiversity | multiple | key biodiversity areas, ecologically intact areas, protected areas |
| 3 | Allan et al | 2017 | Temporally inter-comparable maps of terrestrial wilderness and the Last of the Wild | single | remaining wilderness |
| 4 | Belote et al | 2020 | Mammal species composition reveals new insights into Earth's remaining wilderness | multiple | intactness mammal communities, human footprint |
| 5 | Beyer et al | 2019 | Substantial losses in ecoregion intactness highlight urgency of globally coordinated action | single | habitat intactness |
| 6 | Brooks et al | 2004 | Coverage provided by the global protected area system: Is it enough? | single | species richness, threatened species, protection coverage |
| 7 | Brooks et al | 2006 | Global biodiversity conservation priorities | multiple | high biodiversity threat, low biodiversity threat |

| | | | | | |
|---|---|---|---|---|---|
| 8 | Brum et al | 2017 | Global priorities for conservation across multiple dimensions of mammalian diversity | multiple | mammal phylogenetic diversity, mammal functional diversity, mammal trait diversity |
| 9 | Buchanan et al | 2011 | Identifying priority areas for conservation: A global assessment for forest-dependent birds | single | contribution to forest bird distribution |
| 10 | Buhlmann et al | 2009 | A Global Analysis of Tortoise and Freshwater Turtle Distributions with Identification of Priority Conservation Areas | single | turtle and tortoise richness, turtle and tortoise protection |
| 11 | Butchart et al | 2015 | Shortfalls and Solutions for Meeting National and Global Conservation Area Targets | multiple | species coverage, ecosystem coverage, key biodiversity areas, gross domestic product |
| 12 | Cantu-Salazar et al | 2013 | The performance of the global protected area system in capturing vertebrate geographic ranges | single | richness of under protected vertebrates |
| 13 | Cardillo et al | 2006 | Latent extinction risk and the future battlegrounds of mammal conservation | single | richness latent extinction risk in mammals |
| 14 | Carrara et al | 2017 | Towards biodiversity hotspots effective for conserving mammals with small geographic ranges | multiple | richness range restricted species, richness range restricted evolutionary diversity, richness range restricted threatened species |
| 15 | Ceballos | 2006 | Global mammal distributions biodiversity hotspots and conservation | multiple | mammal species richness, mammal endemic species richness, mammal threatened species richness |
| 16 | Chen and Peng | 2017 | Evidence and mapping of extinction debts for global forest-dwelling reptiles, amphibians and mammals | multiple | extinction depth mammals, amphibians and reptiles, extinction risk mammals, amphibians and reptiles, richness mammals amphibians and reptiles |
| 17 | Cimatti et al | 2021 | Identifying science policy consensus regions | multiple | 63 different conservation priority maps |

| | | | | | |
|---|---|---|---|---|---|
| 18 | Daru et al | 2019 | Spatial overlaps between the global protected areas network and terrestrial hotspots of evolutionary diversity | multiple | phylogenetic diversity, phylogenetic endemism, EDGE |
| 19 | Di Marco et al | 2019 | Wilderness areas halve the extinction risk of terrestrial biodiversity | multiple | vertebrate persistence, plant persistence, wilderness |
| 20 | DiMarco et al | 2012 | A novel approach for global mammal extinction risk reduction | single | extinction risk reduction opportunity |
| 21 | Dinerstein | 2020 | A global safety net to reverse biodiversity loss and stabilize earth climate | multiple | species rarity, distinct species assemblages, rare phenomena, carbon storage, wildlife corridors |
| 22 | Freudenberger et al | 2013 | Nature conservation Priority-setting needs a global change | multiple, weighted | 16 variables including carbon storage, vegetation density, species richness vascular plants, functional richness, forest cover loss and human footprint |
| 23 | Funk et al | 2010 | Ecoregion prioritization suggests an armoury not a silver bullet for conservation planning | multiple | species richness, endemism, endangerment and threat, ecoregions |
| 24 | Giardello et al | 2019 | Global synergies and trade-offs between multiple dimensions of biodiversity and ecosystem services | multiple, weighted | taxonomic, phylogenetic and functional diversity birds and mammals, carbon sequestration, pollination potential and groundwater recharge |
| 25 | Goldstein et al | 2020 | Protecting irrecoverable carbon in Earth's ecosystems | single | manageable carbon, vulnerable carbon, irrecoverable carbon |
| 26 | Gonzales Souza | 2020 | Habitat loss extinction and conservation effort in terrestrial ecoregions | multiple | projected extinction risk, protected area coverage |
| 27 | Grenyer et al | 2006 | Global distribution and conservation of rare and threatened vertebrates | | Bird, mammal and amphibian species richness, endemic richness and threatened richness |
| 28 | Gumbs et al | 2020 | Global priorities for conservation of reptilian PD in the face of human impacts | multiple | reptile phylogenetic endemism, human impact |

| | | | | | |
|---|---|---|---|---|---|
| 29 | Hanson et al | 2020 | Global Conservation of species niches | single | species niches |
| 30 | Hidasi Neto et al | 2015 | Global and local evolutionary and ecological distinctiveness of terrestrial mammals | multiple | ecological and evolutionary distinctiveness mammals, threat status |
| 31 | Hoekstra et al | 2004 | Confronting a biome crisis Global disparities of habitat loss and protection | single | habitat conversion, habitat protection |
| 32 | Howard et al | 2020 | A global assessment of the drivers of threatened terrestrial species richness | single | threatened species richness |
| 33 | Jenkins et al | 2013 | Global patterns of terrestrial vertebrate diversity | multiple | vertebrate richness, endemism and threat |
| 34 | Jetz et al | 2014 | Global Distribution and Conservation of Evolutionary Distinctness in Birds | single | evolutionary distinctiveness birds, threat status, protection coverage |
| 35 | Jung et al | 2021 | Areas of global importance for conserving terrestrial biodiversity, carbon and water | multiple | carbon stock, threatened species, water quality |
| 36 | Kier et al | 2009 | A global assessment of endemism and species richness across island and mainland regions | single | bird, mammal. Amphibian, reptile and vascular plant species richness and endemism |
| 37 | Kullberg et al | 2018 | Using KBAs to guide effective expansion of the global PA network | single | protection coverage threatened vertebrates, key biodiversity areas |
| 38 | Lamoreux | 2006 | Global tests of biodiversity concordance and the importance of endemism | single | Bird, mammal, amphibian and bird species richness and endemism |
| 39 | Loiseau | 2020 | Global distribution and conservation status of ecologically rare mammal and bird species | single | Species richness, ecologically rare species richness, Threatened species |
| 40 | Mazel et al | 2014 | Multifaceted diversity area relationships reveal global hotspots of mammalian species trait and lineage diversity | multiple | mammal species richness, mammal functional diversity, mammal phylogenetic diversity |
| 41 | McDonald et al | 2018 | Conservation priorities to protect vertebrate endemics from global urban expansion | multiple | vertebrate endemism, current land cover, urban expansion |
| 42 | Meyers et al | 2000 | Biodiversity hotspots for conservation priorities | multiple | endemic species richness, habitat loss |
| 43 | Mittermeier et al | 2003 | Wilderness and biodiversity conservation | multiple | wilderness, species richness vascular |

| | | | | | |
|---|---|---|---|---|---|
| | | | | | plants, species richness terrestrial vertebrates |
| 44 | Mittermeier et al | 2015 | Global hotspots for turtle conservation | multiple | species richness tortoises and freshwater turtles, species endemism tortoises and freshwater turtles |
| 45 | Mokany et al | 2020 | Reconciling global priorities for conserving biodiversity habitat | single | assemblage intactness, human footprint |
| 46 | Moran et al | 2017 | Identifying species threat from global supply chains | single | biodiversity footprint, threatened species richness |
| 47 | Naidoo et al | 2008 | Global mapping of ecosystem services and conservation priorities | multiple | carbon sequestration, carbon storage, freshwater provision, grassland production of livestock |
| 48 | Olson et al | 2002 | The global 200 | multiple | species richness, endemic species richness, unusual higher taxa, unusual ecological phenomena, evolutionary phenomena, habitat rarity |
| 49 | Orme et al | 2005 | Global hotspots of species richness are not congruent with endemism or threat | multiple | bird species richness, bird species endemism, bird threatened species richness |
| 50 | Pelletier et al | 2018 | Predicting plant conservation priorities | single | threatened plant species richness |
| 51 | Pollock et al | 2017 | Large conservation gains possible for global biodiversity facets | multiple | mammal and bird species richness, phylogenetic diversity and functional diversity |
| 52 | Pouzols et al | 2014 | Global PA expansion is compromised by projected land-use and parochialism | multiple | species richness, current protection coverage, projected future land-use |
| 53 | Riggio et al | 2020 | Global human influence maps reveal clear opportunities in conserving Earths remaining intact terrestrial ecosystems | single | anthromes, human footprint, Low impact areas, Global human modification |
| 54 | Rodriguez et al | 2004 | Global gap analysis Priority regions for expanding the global protected-area network | multiple | species richness mammals, amphibians, freshwater turtles, tortoises and |

| | | | | | |
|---|---|---|---|---|---|
| | | | | | threatened birds, protected areas |
| 55 | Roll et al | 2017 | The global distribution of tetrapods reveals a need for targeted reptile conservation | single | species richness reptiles, species richness terrestrial vertebrates |
| 56 | Rosauer et al | 2017 | Phylogenetically informed spatial planning is required to conserve the mammalian tree of life | single | mammalian phylogenetic diversity, species richness |
| 57 | Safi et al | 2013 | Global Patterns of Evolutionary Distinct and Globally Endangered Amphibians and Mammals | multiple | evolutionary distinctiveness amphibians and mammals, threat status amphibians and mammals, |
| 58 | Schipper et al | 2008 | The status of the worlds land and marine mammals: Diversity, threat and knowledge | multiple | terrestrial and marine mammal species richness, endemic species, threatened species and phylogenetic diversity |
| 59 | Soto Navarro et al | 2020 | Global biodiversity conservation priorities | multiple | species richness–area of habitat, rarity-weighted richness–area of habitat, mean species abundance, biodiversity intactness index, biodiversity habitat index, above- and below ground terrestrial carbon storage, |
| 60 | Stuart et al | 2004 | Status and Trends of Amphibian Declines and Extinctions Worldwide | multiple | species richness, declining species, enigmatic species, habitat loss, over exploitation |
| 61 | Veach et al | 2017 | Species richness as criterion for global conservation area placement leads to large losses in coverage of biodiversity | multiple | species richness vertebrates, threatened species richness vertebrates, endemic species richness vertebrates |
| 62 | Venter et al | 2014 | Targeting global protected area expansion for imperilled biodiversity | multiple | area protected, opportunity cost, number of species protected |

| | | | | | |
|---|---|---|---|---|---|
| 63 | Voskamp et al | 2017 | Global patterns in the divergence between phylogenetic diversity and species richness in terrestrial birds | single | species richness birds, phylogenetic diversity birds |
| 64 | Watson et al | 2018 | Protect the last of the wild | single | remaining wilderness |
| 65 | Yang et al | 2020 | Cost effective priorities for the expansion of global terrestrial protected areas Setting post 2020 global and national targets | multiple | crisis ecoregions, biodiversity hotspots, endemic bird areas, key biodiversity areas, centres of plant diversity, global 200s, and intact forest landscapes, human footprint, human modification, low human impact areas |

### 2. Supplementary methods and materials

Below we provide a detailed description of the protected area datasets and the individual indicators underlying the conservation objectives and how these data were derived.

#### 2.1 The protected area dataset

The potential sites currently included in the analysis are either included as protected areas, IUCN category I or II, or listed as a Natural World Heritage Site (WHS), or registered as a Key Biodiversity Area (KBA). The shapefiles for the IUCN protected areas and the Natural World Heritage Sites (WHS) were derived from the World Database on Protected Areas [1] excluding those sites for which only point data was available. The shapefiles for the Key Biodiversity Areas (KBAs) were obtained from BirdLife International [2].

There are various sites in the world where the WHS sites or the KBAs overlap with the IUCN protected areas. We resolved all such spatial conflicts by retaining the shapefile with the higher protection status where different shapefiles overlapped (IUCN > KBA > WHS). For example, WHS sites that were embedded within an IUCN protected area as well as KBAs that overlapped with an IUCN protected area were excluded from the analysis. In some instances, there was only a partial overlap of either a KBA or WHS site with an IUCN protected area or a KBA overlapped with an IUCN protected area but was considerably larger (Fig S1). For these cases we kept both shapefiles in the analysis. This was the case for 17 sites (Table S2).

We sampled all protected area polygons into a grid of 0.5° longitude x 0.5° latitude, deriving the percentage overlap of each polygon with the grid cells.

To estimate the potential impacts of projected land-use change around the protected areas, we derived 50 km buffers around each protected area polygon and then sampled these into the grid as described above.

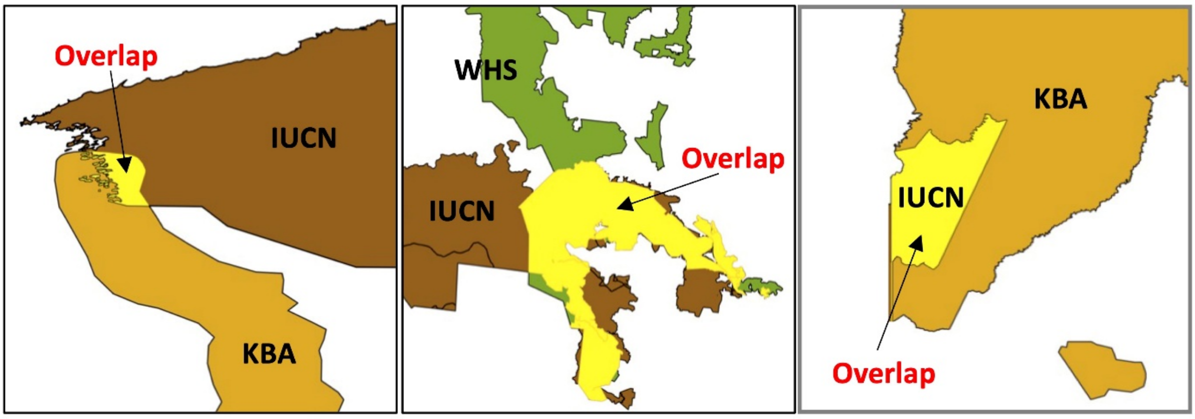

**Fig. S1:** Examples of marginal, partial and full overlap of two shapefiles. Left shows a KBA (orange) that has marginal overlap with an IUCN site (brown). Centre shows a WHS site (green) that partially overlaps with an IUCN site but is kept because it is considerably larger than the area already covered by the IUCN site. Right shows an IUCN site that is embedded within a KBA, here too the KBA is kept because it is considerably larger than the IUCN site.

**Table S2:** Number of sites that had partial, marginal or full overlap with another site included in the dataset.

| Overlapping sites | Type of overlap | Number of occasions |
|---|---|---|
| IUCN + KBA | marginal | 8 |
| IUCN + KBA | partial | 2 |
| IUCN + KBA | embedded | 1 |
| IUCN + WHS | marginal | 3 |
| IUCN + WHS | embedded | 1 |
| WHS + KBA | marginal | 2 |
| **Total** | **17** (1.3% of sites included) | |

#### 2.2 The conservation objectives

The six different conservation objectives which are included in the decision support tool are biodiversity, wilderness, climatic stability, land-use stability, climate protection and size. Each of these objectives consists of one or several underlying biogeographic indicators. The detailed description which variables are included in each of the conservation objectives and how these variables were derived is given below.

##### *2.2.1 Biodiversity*

The biodiversity objective includes three different variables: the species richness of the site, the average degree of endemism across the species occurring within the site, and the evolutionary diversity of the species occurring in the site.

*Species richness (SR)*

The species richness (Fig. S2) for four taxa of terrestrial vertebrates was derived from BirdLife (birds), IUCN (mammals, amphibians) or GARD (reptiles) range-map polygons, which were gridded to the 0.5°

grid [3–5]. The species ranges were stacked to obtain species lists for each grid cell. The resulting species matrix was then merged with the site grid and the unique species across all grid cells within each site grid were summed up as the SR value for the site. For the site selection, sites with a high SR are of high value, whereas sites with a low SR are of less value.

*Species endemism: corrected range size rarity (RSR)*

To capture unique biodiversity, we included a measure for the number of range-restricted (endemic) species within a protected area, the so-called range size rarity (RSR, Fig. S3) which has been used as a proxy for species endemism [6]. This is derived by summing the species for each grid cell, including weights that reflect species' range sizes. Usually range size rarity is calculated by weighting each species by the inverse of its range extent (e.g. number of cells occupied globally), so that species within a given grid cell have larger weights if they occur in very few other grid cells [7,8]. The resulting values are highly correlated to species richness, because the weighted species values are summed up per grid cell [6]. Therefore, we corrected for species richness by dividing the weighted range size rarity value by the total number of species within the grid cell following Crisp *et al.* 2001. Using this corrected range size rarity (RSR) as a measure instead of the raw number of endemic species is of advantage because there is no arbitrary cut off to define endemic species. Whereas endemism is often calculated based on the 25% of the species with the smallest range size in the world, range size rarity is based on a gradient of how endemic species are on average within a site.

Site specific RSR values were derived for the four vertebrate taxa in the same way as SR values, by merging the species matrix (containing the species-specific range size rarity values for each grid cell) to the site grid. summing the RSR values of the unique species across all grid cells of the site. For the site selection, sites with a high RSR are of high value, whereas sites with a low RSR are of less value.

*Evolutionary diversity: phylogenetic endemism (PE)*

Evolutionary diversity was included to evaluate how evolutionarily unique the species within a protected area are. Measures of phylogenetic diversity, as Faith PD, can give an idea of how much evolutionary history is stored within a set of species [9]. A high amount of evolutionary history has been linked to higher productivity and stability of ecosystems [10,11]. Evolutionary diversity was calculated using phylogenetic endemism (PE), which is a combined measure of phylogenetic diversity and uniqueness of a species community [12]. PE (Fig. S4) identifies areas with high numbers of evolutionarily isolated and geographically restricted species. Additionally to the summed shared evolutionary history of a species assemblage, PE therefore incorporates the spatial restriction of phylogenetic branches covered by the assemblage [12]. PE was calculated following the method developed by Rosauer *et al* (2009). To derive the PE values, we used the phylogenetic supertree for all four terrestrial vertebrate taxa from Hedges *et al.* [13], which was combined with the aforementioned species range-map data from IUCN and BirdLife International [14]. The number of species for which both distribution and phylogenetic data were available differed across taxa, but all analyses included high percentages of the globally known species in each taxon (Table S3). PE was derived for each 0.5° grid cell and then the PE for each protected area was calculated as mean PE across all grid cells within the area polygon. For the site selection, sites with a high PE are of high value, whereas sites with a low PE are of less value.

**Table S3:** The number of species in each class of terrestrial vertebrates for which phylogenetic data was available, and the number of species that were included in the analyses for species richness and endemism but which are missing in the phylogenetic endemism analysis. We also give the total number of species with distribution data and the corresponding percentage of known species represented in each taxon, following the respective taxonomy [3–5].

| Taxa | Species w. phylogenetic + distribution data | Species w. distribution data only | Total | % |
|---|---|---|---|---|
| Birds | 8296 | 1360 | 9656 | 86 |
| Mammals | 4867 | 113 | 4980 | 98 |
| Amphibians | 6051 | 145 | 6196 | 98 |
| Reptiles | 8801 | 1263 | 10064 | 87 |

2.2.2 *Ecosystem intactness*

The ecosystem intactness objective includes three different variables: biodiversity intactness index, human footprint and recent land-use change.

*Biodiversity intactness index (BII)*

The biodiversity intactness index represents the modelled average abundance of present species, relative to the abundance of these species in an intact ecosystem [15]. This means it gives an indication how much species abundances in an area have already changed due to anthropogenic impacts such as land-use change. We used the global map of the Biodiversity Intactness Index (BII) provided by Newbold et al 2018 (see [16] for a detailed description of how the BII is derived). The values were extracted for each grid cell, grid cell values were weighted by their percentage overlap with the protected area polygon, and then weighted mean BII values were derived for each protected area. For the site selection, sites with a low BII within the protected area are of lower value, whereas sites with a high BII are of higher value.

*Human footprint (HFP)*

As a measure of how pristine the protected areas still are in general, a measure of the human footprint within the area was included. Estimates of the human footprint (HFP) within protected areas were derived using the data of Venter *et al.* 2016 [17]. We used the standardised HFP that was provided by Venter et al. and includes data on the extent of built environments, cropland, pasture land, human population density, night-time lights, and the density of railways, roads and navigable waterways. We aggregated the HFP layers to the half degree resolution, derived HFP values for each grid cell, weighted grid cells by their percentage overlap with the protected area polygon and derived the mean HFP for each protected area. For the site selection, sites with a high human footprint within the protected area are of lower value, whereas sites with a low human footprint are of higher value.

*Land-use change*

To derive past changes in the land cover of the protected area we calculated the average percentage of the site altered from biomes (natural land cover classes) to human dominated land cover classes (anthromes; i.e., urban/semi-urban areas and cultivated areas). The time series of fractions of land cover classes, ranging from 1992 – 2018, was obtained from the GEOEssential project [18]. The land cover classes used in this were derived from the ESA CCI Land Cover and were available on a 30km grid. We calculated the total percentage change from biomes to anthromes between the years 1992 and 2018 and aggregated the data into the half degree grid. The summed changes for each protected area polygon were derived from the grid cell values weighted by the percentage overlap of grid cells and polygon. For the site selection, sites with a high percentage land-use change between 1992 and 2018 are of lower value and sites with a low percentage land-use change are of higher value.

##### *2.2.3 Climatic stability*

The climatic stability objective consists of two different variables: the climatic stability of biodiversity using the four terrestrial vertebrate taxa, and the projected tree cover change.

*Climatic stability of biodiversity*

To assess the climatic stability of a protected area, we evaluated the potential impacts of climate change on the biodiversity within the site. Climate change is already driving observable shifts in species distributions and it is well known that many taxa are shifting their ranges towards higher latitudes [19,20]. However, idiosyncratic species responses to climate change have also been observed [21–23]. These range shifts have the potential to reshuffle species assemblages, which can have highly unpredictable impacts on the assemblage (e.g., changes in prey-predator balance or competition). We assume that species assemblages which are predicted to change only weakly in composition in the future or to experience very few species losses are under less risk from climate change than species assemblages projected to experience a lot of reshuffling. Under this assumption, we defined the inverse of projected turnover in species as an indicator for climatic stability, and calculated climatic stability for each protected area until 2050. The projected turnover is calculated for each of the four vertebrate taxa based on species-level range-map projections derived from species distribution models (SDMs). The SDMs have been published previously (see [24] for a detailed account of the modelling methods) and are based on an ensemble of two modelling algorithms (Generalized additive models and Generalized Boosted Regression Models) and four different Global Climate Models (GCMs; MIROC5, GFDL-ESM2M, HadGEM2-ES and IPSL-CM5A-LR). These models use the meteorological forcing dataset EartH2Observe, WFDEI and ERA-Interim data, which were merged and bias-corrected for ISIMIP

(EWEMBI [25]), as dataset for the current climatic conditions (from 1980 – 2009). As future climate dataset, they rely on bias-corrected global climate scenarios produced by ISIMIP phase 2b [26]. Here we used the projections assuming a medium dispersal scenario (allowing dispersal across a distance equal to half the largest radius of the range polygons of a species), and a medium concentration pathway (RCP 6.0). Species with range extents of fewer than 10 grid cells were excluded from the modelling. In total we had modelled distributions available for 22,652 vertebrate species (see Table S4) on the 0.5° grid. To derive species lists per site we applied species-specific thresholds that maximized the fit to the current data, using the true skill statistic (MaxTSS), to translate the projected probabilities of occurrence into binary presence absence data [27]. For each site, all species that were projected to occur currently and/or in future (2050) were extracted. Turnover was then calculated between the current and future species assemblage of a site, using the formula for Bray Curtis dissimilarity [28]:

$$B_{ij} = \frac{2C_{ij}}{S_i + S_j}$$

Where $S_i$ and $S_j$ are the species counts at the two points in time, and $C_{ij}$ are the counts of species found in both sites. For the site selection, sites with a high projected turnover as a consequence of global climate change are of low value, whereas sites with a low projected turnover are of high value.

| Taxa | Species with SDM | Species without SDM | Total | % |
|---|---|---|---|---|
| Terrestrial birds | 8986 | 896 | 9882 | 91 |
| Terrestrial mammals | 4307 | 968 | 5275 | 82 |
| Amphibians | 3063 | 3317 | 6380 | 48 |
| Reptiles | 6296 | 3768 | 10,064 | 60 |

**Table S4:** The number of species in each class of terrestrial vertebrates for which species distribution models could be built and which were included in the analyses for climate stability of biodiversity. The total species number is the number of species with range maps, we also give the corresponding percentage of species with range maps models could be built for (cf. Table S3).

*Projected tree cover change*

We included the projected potential forest cover change from 1995 until 2050 based on the projected change in tree cover of the LPJ-GUESS process-based dynamic vegetation-terrestrial ecosystem model [29]. This variable captures changes in forest cover but not necessarily changes in other vegetation types, e.g. the desertification of grasslands and drylands. The projected changes in forest cover are driven by climate and CO2 changes but do not include projected changes in land-use. The climate input for the model was derived from the ISIMIP2b simulations (see detailed description above under Climatic stability of biodiversity). The projected change in tree cover was provided as a percentage per grid cell. The grid cell values were weighted by their percentage overlap with the protected area polygon, and then the weighted mean percentage change in tree cover was derived for each protected area. Both a strong decrease as well as a strong increase in tree cover could equal a risk for a site, e.g. a projected loss in tree cover could be a risk for a forest whilst a projected increase could be a risk for grasslands.

Therefore, sites with a low projected change in tree cover, in either direction, are of higher value, for the site selection, whereas sites with a high projected change in tree cover are of lower value.

##### 2.2.4 *Land-use stability*

To assess the potential impacts projected future land use change we used predictions of the change in pastures, croplands and biofuel croplands around the sites.

*Projected land-use change around the site*

Projected land–use change was derived from the ISIMIP2b simulations of current and future land-use for 1995 and 2050, based on the MAgPIE and REMIND-MAgPIE model [30–32], using the assumptions of population growth and economic development as described in [26]. Land-use change models accounted for climate impacts (e.g., on crop yields) and were driven with the same climate model projections as the SDMs used to derive climatic stability (see above). The ISIMIP land-use scenarios provide percentage cover of six different land-use types (urban areas, rainfed crop, irrigated crop, pastures, as well as rainfed and irrigated bioenergy crops) at a spatial resolution of 0.5°. We averaged the land-use change for each land-use type across the four GCMs. We then calculated a summed value of land-use change (cropland, biofuel cropland and pastures) between the two different time periods (1995 and 2050), per grid cell. To get an estimate of the potential pressure that future land-use change could put on a protected area, we derived the mean and maximum values of the projected land-use change across all grid cells in the 50 km buffer zone around each protected area (see section 1.1 above). The grid cell values were weighted by their extent of overlap with the buffer zone to derive the final value for each site. For the site selection, sites with a high projected land-use change around the protected area are of low value, whereas sites with a low projected land-use change are of higher value.

##### 2.2.5 *Climate protection*

We used data on carbon stored in vegetation and soils as an indicator of the potential of a site to contribute to climate protection. The climate protection objective includes three different indicators, the amount of manageable carbon stored in the site, the amount of vulnerable carbon and the amount of irrecoverable carbon.

*Manageable carbon*

Here we used the estimated amount of manageable carbon as provided by Noon *et al* 2021. Manageable carbon is defined by Goldstein *et al* 2020, as an ecosystems carbon stock that is primarily affected by human activities that either maintain, increase or decrease its size. This layer is derived from a comprehensive suite of carbon datasets across terrestrial, coastal and freshwater ecosystems globally. It includes the amount of carbon stored in the above and below ground vegetation as well as soil organic carbon stocks up to 30 cm depth, or up to 100 cm within inundated soil, as these depths are most relevant to common disturbances [34]. We aggregated the carbon data[33] to a 0.5° resolution and calculated the amount of manageable carbon storage in t per grid cell. Aggregating the data to the same resolution as the other datasets before using it for the analysis is necessary to speed up data processing for the decision support tool. The grid cell values were weighted by their percentage overlap with the protected area polygon to derive the final mean manageable carbon storage value per site. For the site selection, sites with lower baseline carbon stocks are of lower climate protection value, whereas sites with higher baseline carbon stocks are of higher climate protection value.

*Vulnerable carbon*

Vulnerable carbon is defined by Goldstein *et al* (2020) as the amount of manageable carbon, described above, that is likely to be released through typical land conversion in an ecosystem. Considered conversion drivers here were agriculture for grasslands, peatlands and tropical forests; forestry for boreal and temperate forests; and aquaculture or development for coastal ecosystems [34]. We aggregated the vulnerable carbon data[33] to a 0.5° resolution and calculated the carbon storage in t per grid cell. The grid cell values were weighted by their percentage overlap with the protected area polygon to derive the final mean vulnerable carbon storage value per site. For the site selection, sites with higher vulnerable carbon stocks are allocated a higher suitability for long-term conservation than sites with lower vulnerable carbon stocks.

*Irrecoverable carbon*

Irrecoverable carbon is defined as the amount of the vulnerable carbon, described above, that if it is lost through typical land conversion actions, cannot be recovered over the following 30 years, even if human activities cease [34]. We aggregated the irrecoverable carbon data[33] to a 0.5° resolution and calculated the carbon storage in t per grid cell. The grid cell values were weighted by their percentage overlap with the protected area polygon to derive the final mean irrecoverable carbon storage value per site. For the site selection, sites with higher irrecoverable carbon stocks are allocated a higher suitability for long-term conservation than sites with lower irrecoverable carbon stocks.

2.2.6 *Large size*

For the conservation objective of size, we preselected sites that are larger than 2000 $km^2$. Though being a quite arbitrary threshold, the minimum size was set as a result of the LLF stakeholder debate based on the assumption that larger areas have a higher potential to support populations of target species and to maintain functioning ecosystems in the long term [35,36]. Even for areas above this threshold, the size of the site is still an important criterion under this reasoning, and we used the extent of the site polygon as variable / indicator of this. The Area in $km^2$ was derived from the site polygons (see 1.1 The protected area dataset). The IUCN and World Heritage sites were provided in Mollweide projection. To calculate the $km^2$ extent, the entire dataset was projected to Mollweide projection and $km^2$ were then measured in Q GIS using the area measurement tool [37].

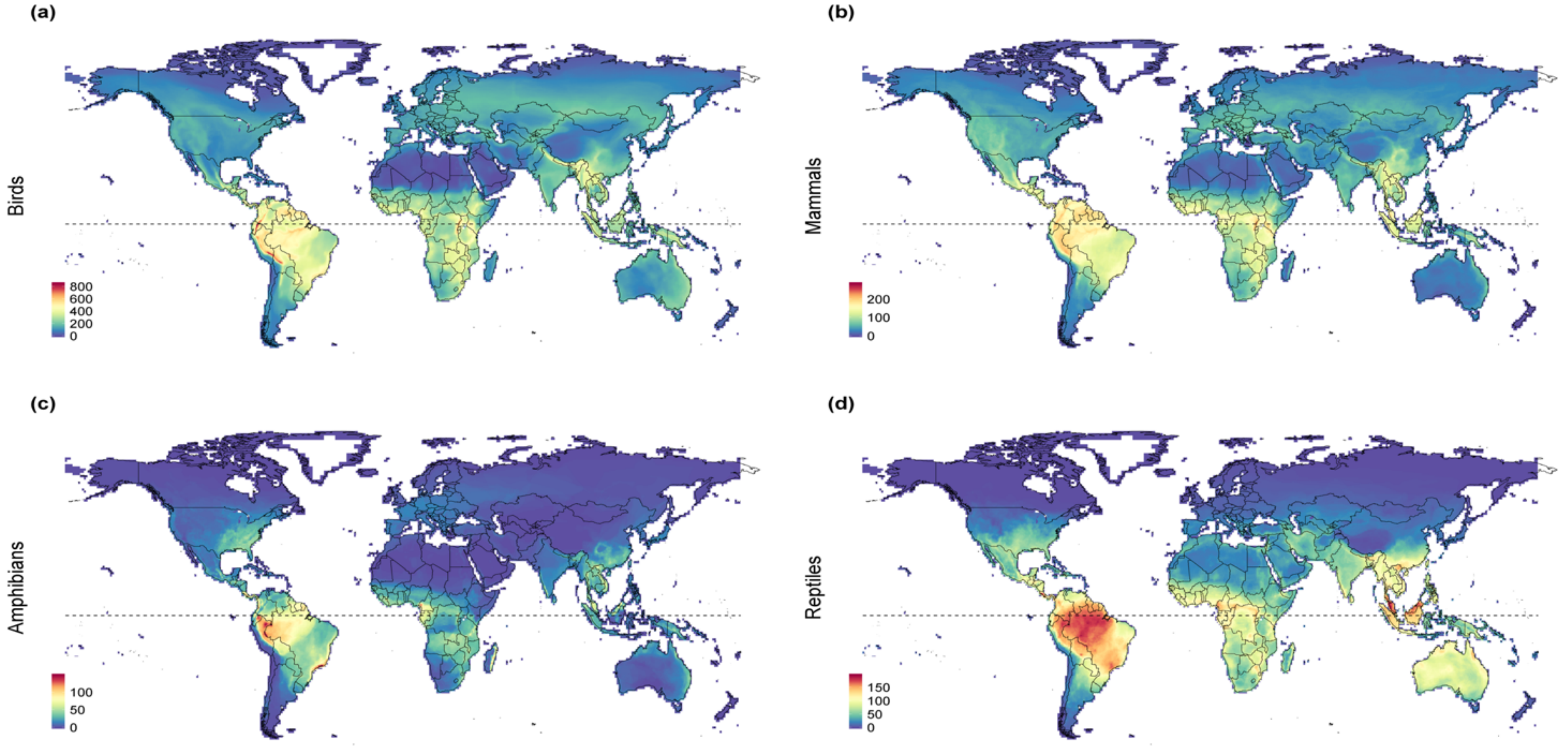

**Fig. S2:** Global species richness for all four taxa of terrestrial vertebrates (birds, mammals, amphibians and reptiles), calculated on a 0.5-degree grid. *Note that the colour scale extent differs between the different taxa.*

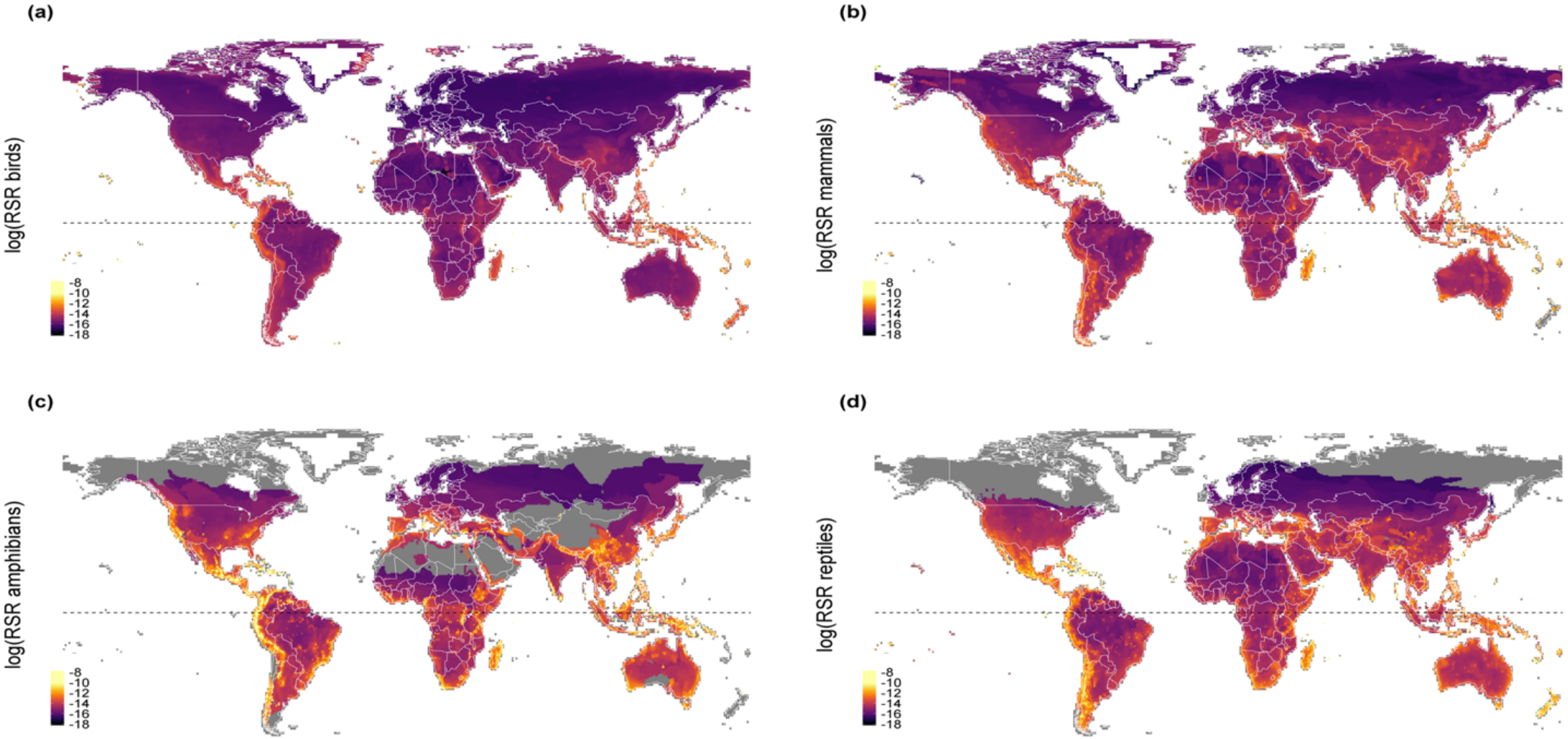

**Fig. S3:** Global corrected range size rarity for all four taxa (birds, mammals, amphibians and reptiles), calculated on a 0.5-degree grid. Corrected range size rarity is the number of species weighted by their inverse range size and divided by the total number of species, shown here on a logarithmic scale. *Note that the scale differs between the different taxa.*

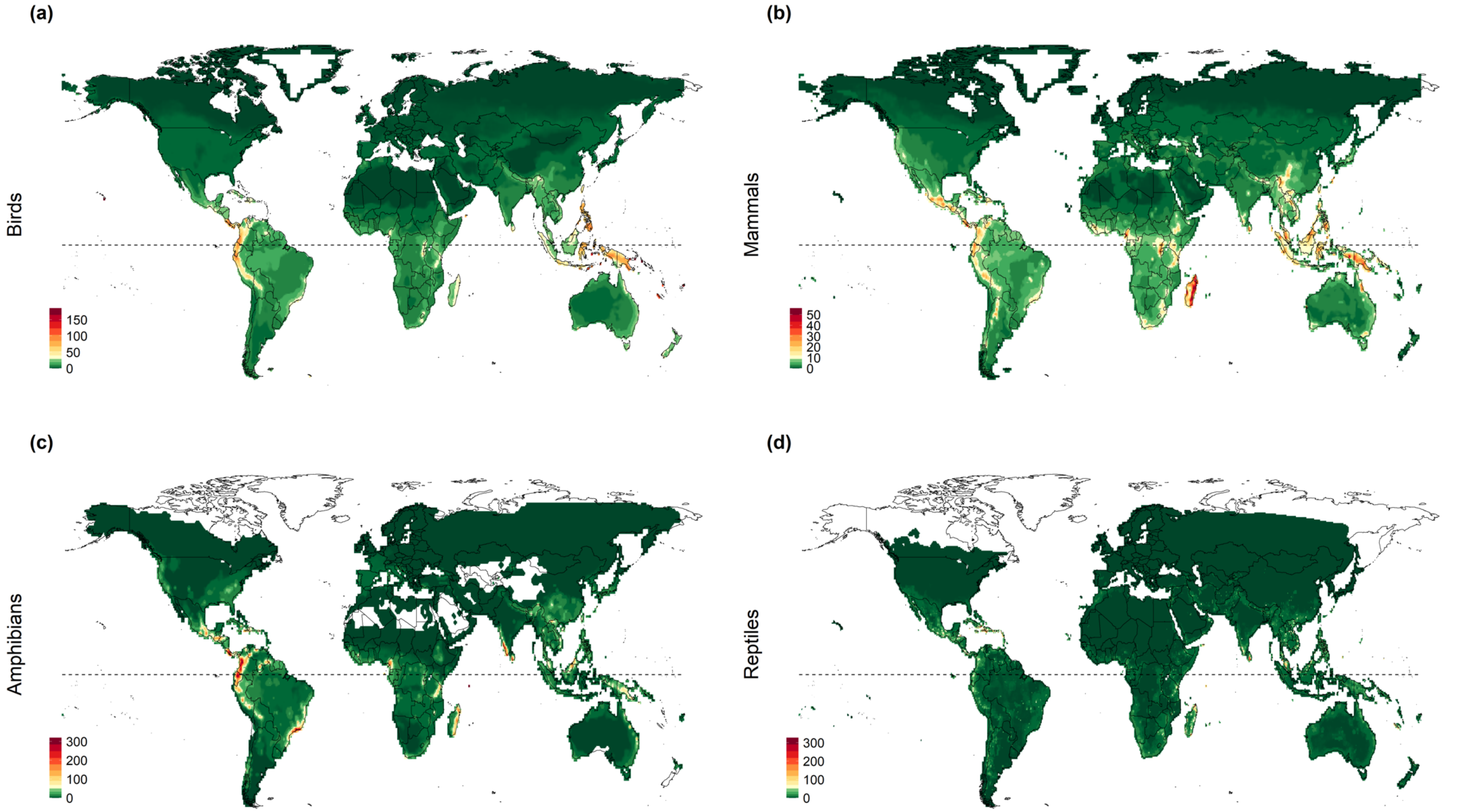

**Fig. S4:** Global patterns of phylogenetic endemism for all four taxa (birds, mammals, amphibians and reptiles), calculated on a 0.5-degree grid. Phylogenetic endemism is calculated by summing the shared evolutionary history of a species assemblage and combining it with information on the range extent of the individual species. *Note that the scale differs between the different taxa.*

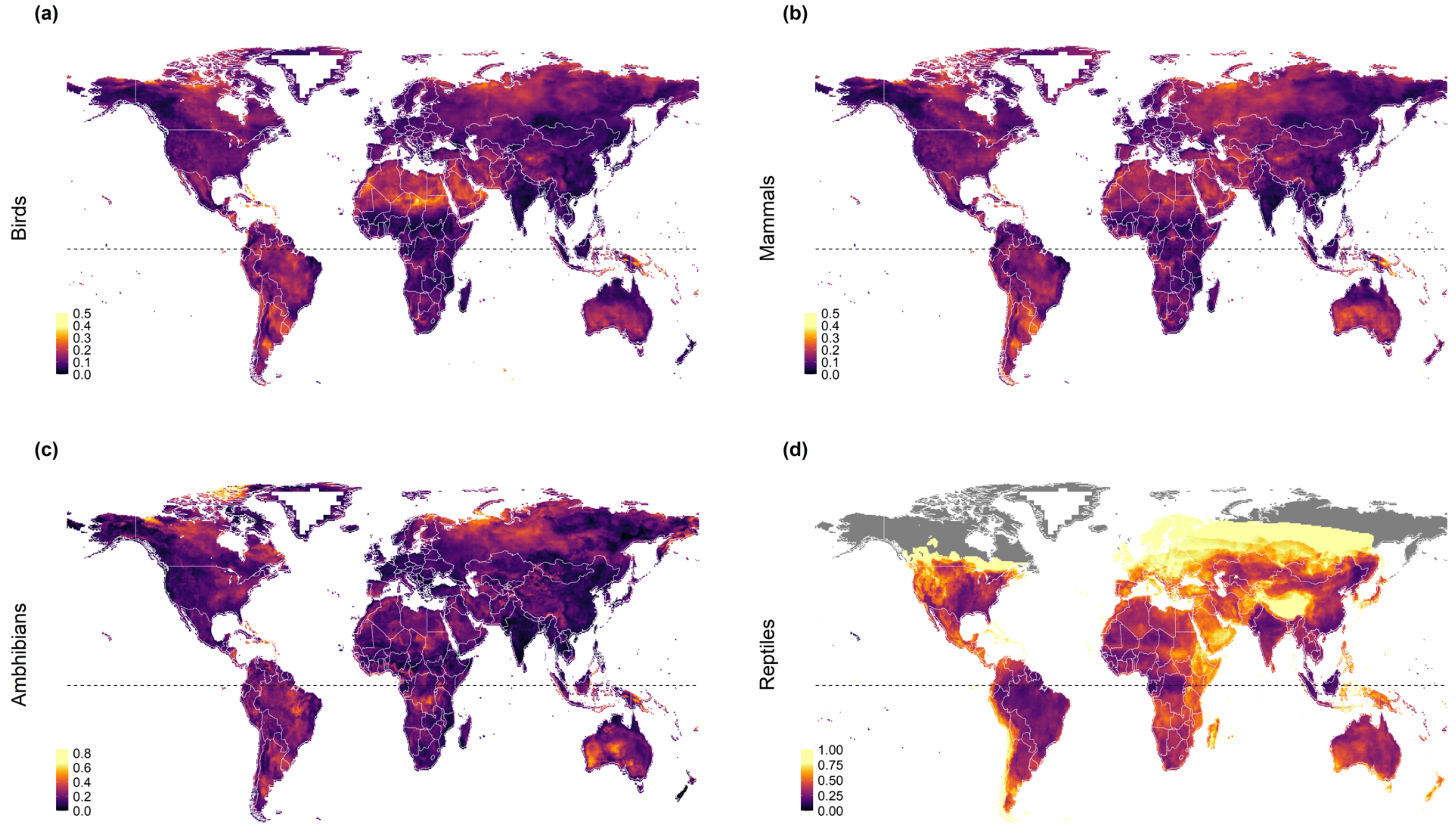

**Fig. S5:** Projected assemblage-level turnover values under climate change for all four taxa (birds, mammals, amphibians and reptiles), calculated on a 0.5-degree grid. Turnover ranges from 0 (low) to 1 (high) and was calculated between the projected current species compositions (1995, average climate projections from 1980 – 2009) and the projected future species compositions (2050, average climate projections 2035 - 2064) under a medium emission scenario (RCP 6.0) and assuming a medium dispersal scenario. *Note that the scale differs between the different taxa.*

#### 2.3 Scaling and weighting the indicators for the site evaluation

We calculated values for each indicator variable for each site included in the conservation decision support tool. For both, summarizing the individual indicators into conservation objectives and weighing them in the decision support tool as well as for the PCA, these values need to be scaled. Therefore, all variables were scaled from 0 to 1, where high values have high priority and low values have low priority for conservation. For some of the variables the original data is opposite to this scale (e.g. for the human footprint an area with a high value is of lower conservation value than a low value); therefore we multiplied this variable by -1 after scaling them. The variables for which the scale was reversed were human footprint, recent land-use change, and land-use stability and climate stability of species communities and tree cover change. For the change in tree cover we assumed that both high positive values (i.e strong increase in tree cover) as well as high negative values (i.e. strong decrease in tree cover) are not desirable. Therefore, we changed the original variable into absolute values. It is interpreted the same way as all other variables with high values (1) being good and low values (0) being less desirable for conservation.

To aggregate indicators that belong to one conservation goal into a single variable, we averaged the scaled variables and rescaled the resulting values to range from 0 to 1.

The three carbon storage variables that are included in the climate protection goal were the only set of variables that are nested (i.e. irrecoverable carbon is part of the vulnerable carbon stock, and vulnerable carbon is part of the baseline carbon stock in the site). We treated the carbon stock variables the same way as the other variables. This is valid under the assumption that the different carbon variables are each of comparable priority. For example, the protection of irrecoverable carbon might arguably be as important for climate protection as the sole protection of manageable carbon. Taking the average across the three variables acknowledges these values. Assume that there are two sites, one with a high amount of manageable carbon but no irrecoverable carbon, and one with lower manageable carbon but a high amount of that being irrecoverable; these sites come out with a similar averaged value. Thus, although the second site has less carbon storage potential in total, some of it is of high importance for climate protection (see correlation matrix for carbon storage Fig. S8).

#### 2.4 The principal component analysis (PCA)

To investigate trade-offs and synergies between the different indicators included in the conservation goals, we used a principal component analysis. The analysis was conducted in R (version 4.1.1), using the "prcomp" function from the "stats" package [38]. All variables were scaled and shifted to be zero centered before the analysis. The PCA plots (Fig 6) were generated using the "fviz_pca" function of the "factoextra" package [39].

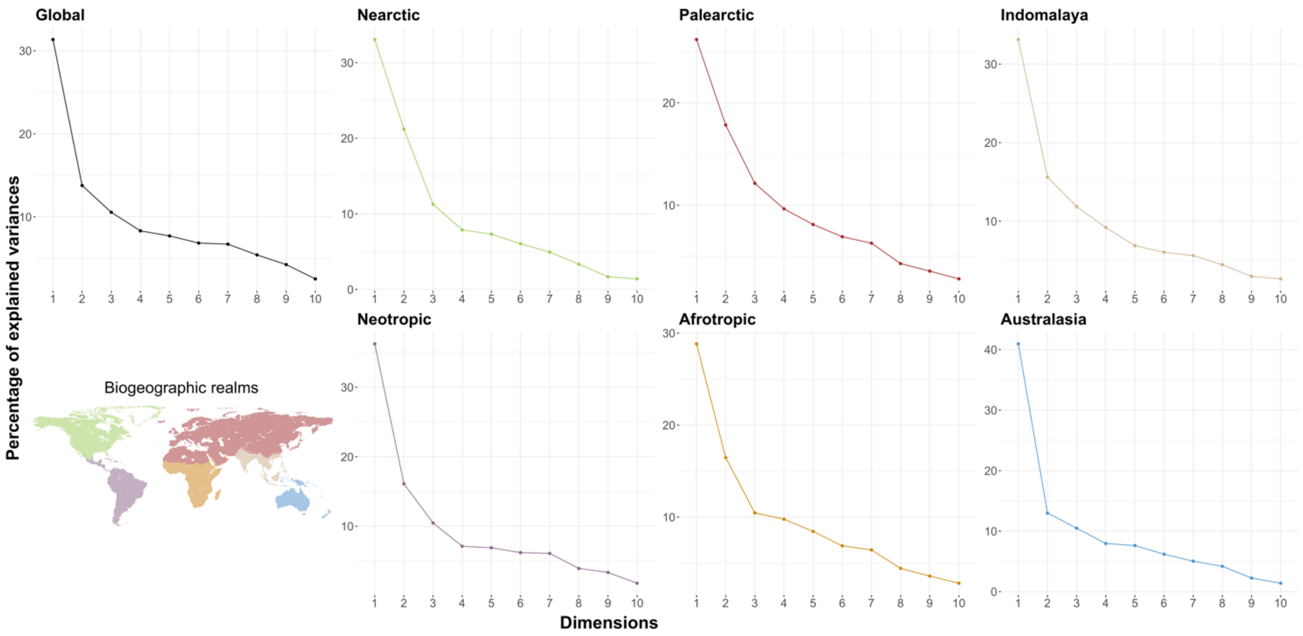

**Fig. S6:** The percentage of variance explained across the different dimensions of the principal components analysis, shown for the global PCA and the realm-wise PCAs.

#### 2.5 Sensitivity of site rankings

We assessed the correlation between the scaled values that were calculated for each conservation objective for each site included in the analysis. As expected, based on the identified synergies and trade-offs in the PCA analysis, the Pearson correlation coefficients between the different conservation objectives were low (Fig. S7). The highest correlation (r=0.58) was found between the Biodiversity and the Climate Protection objective.

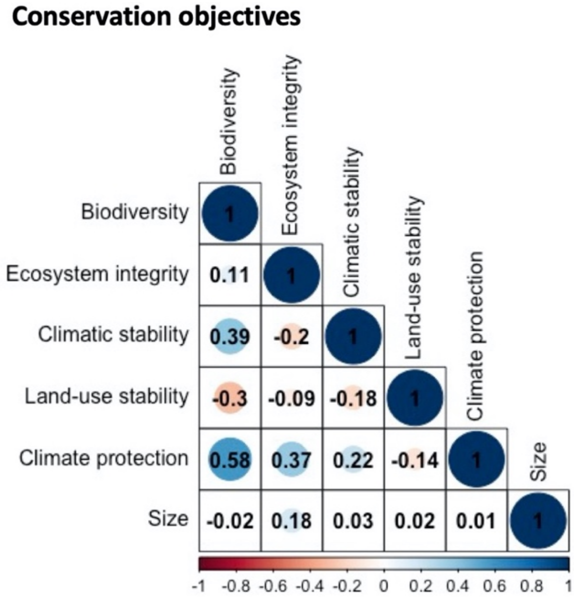

**Fig. S7:** Correlation matrix of the different conservation objectives included in the conservation decision support tool, n=1347.

The correlation between the different indicators included within the conservation objectives varied between the objectives (Fig. S8). Within the biodiversity (Pearson's r >0.20 and <0.77) and the climate protection (Pearson's r >0.85 and <1) objective, the individual indicators tended to be more strongly correlated than within the ecosystem integrity (Pearson's r >0.01 and <0.08) and climatic stability (Pearson's r >-0.08 and < 0.88) objective.

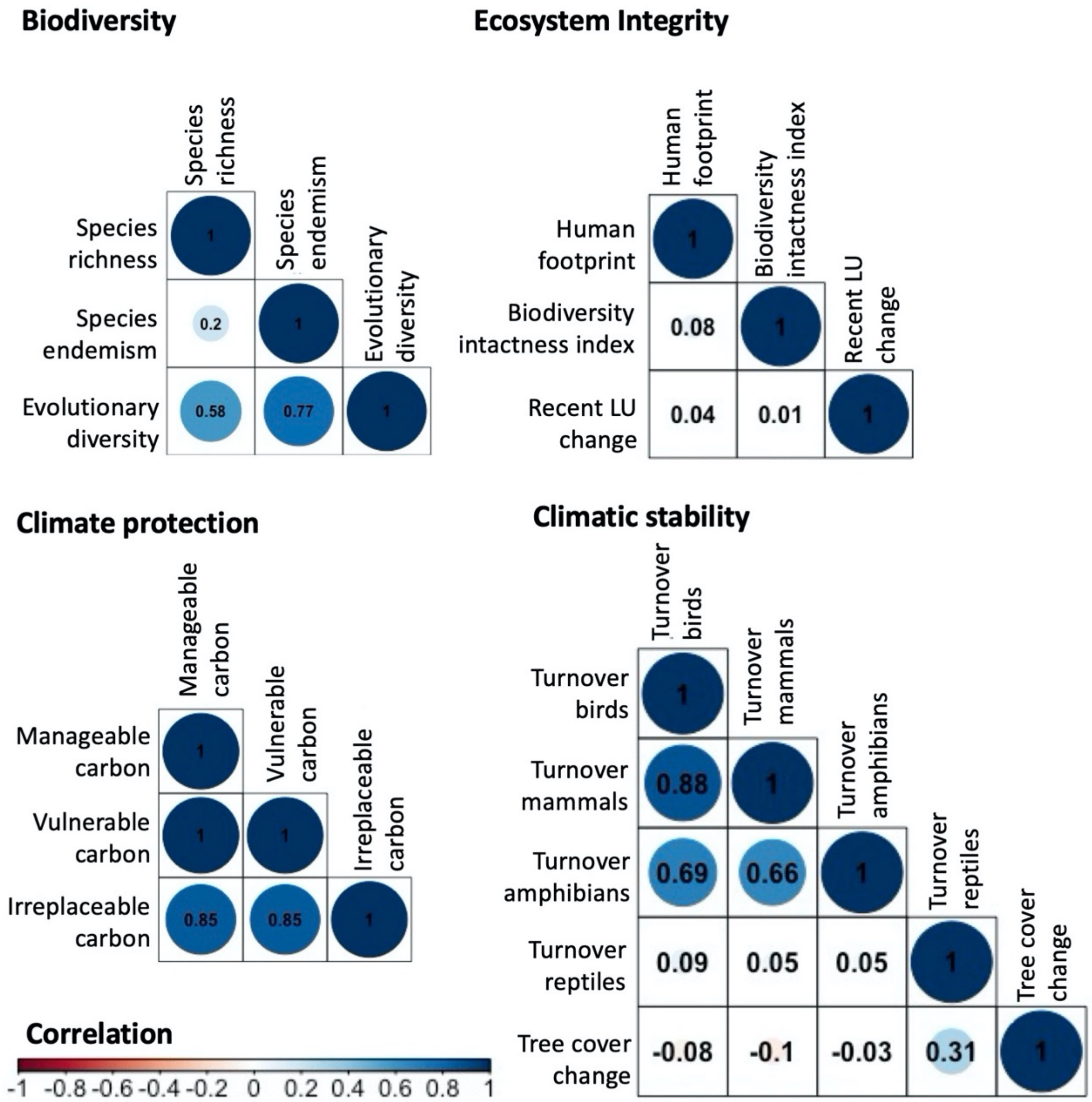

**Fig. S8:** Correlation matrix of the different indicators included in the biodiversity, ecosystem integrity, climate protection and climatic stability, n=1347.

The conservation decision support tool allows the selection and weighting of the individual conservation objectives, but does not offer a sub-weighting of the individual indicators included within an objective. To investigate how much the rankings of individual sites could vary if they were evaluated based on a single indicator instead of the combined objective values, we looked at the changes in rank positions across all sites included in the analysis (Fig. S9 to Fig. S11). For comparison, we also looked at the changes in ranking positions between the conservation objectives, evaluating sites based on one objective at a time. We found that the average range change between the different conservation objectives was 435 rank positions (Fig. S9). Looking at the changes in rank positions within the individual conservation objectives, we found that the magnitude of the average change in rank position differed strongly between the different objectives (Fig S10 and Fig S11). Whilst the average change across the three biodiversity indicators across all sites was 221 rank positions, the average change across the two climatic stability indicators was 377 rank positions. Though there is variation in the ranking positions between the individual indicators included within the conservation objectives, the changes in ranking positions between the conservation objectives is markedly higher.

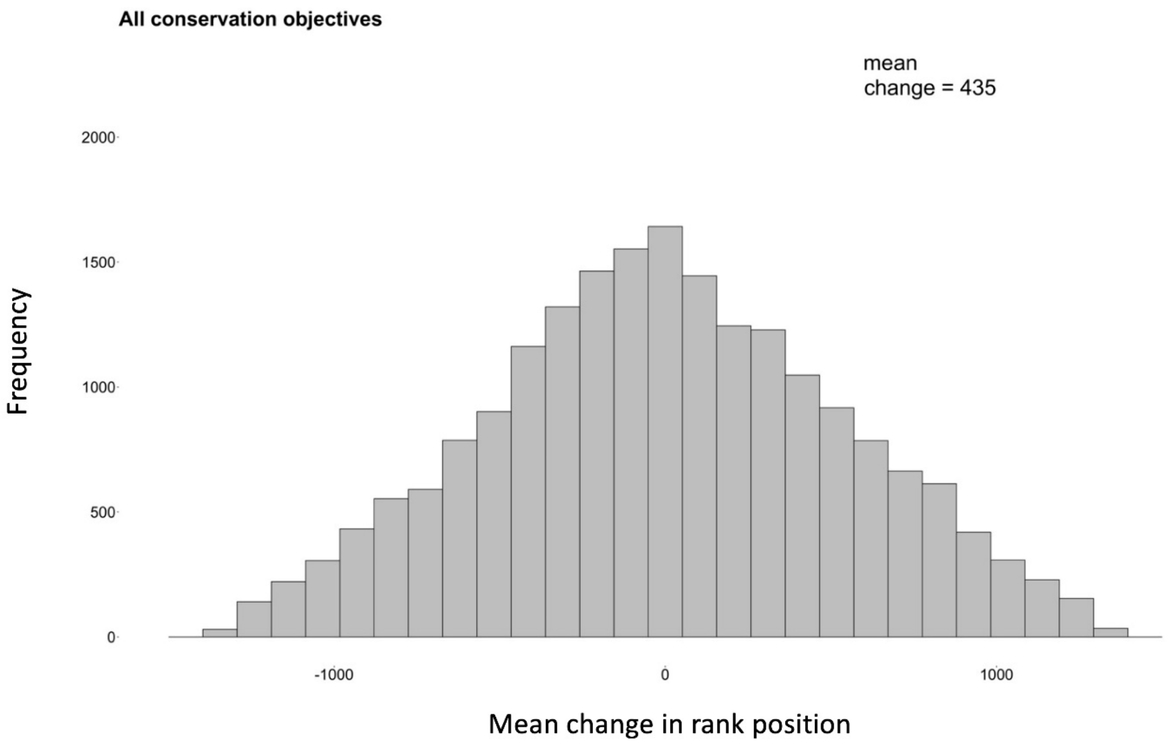

**Fig. S9:** Mean change in rank positions across all sites for the six different conservation objectives. To assess the mean change in rank position, all sites were ranked for each conservation objective individually and the average change in rank position per site was compared across the individual rankings.

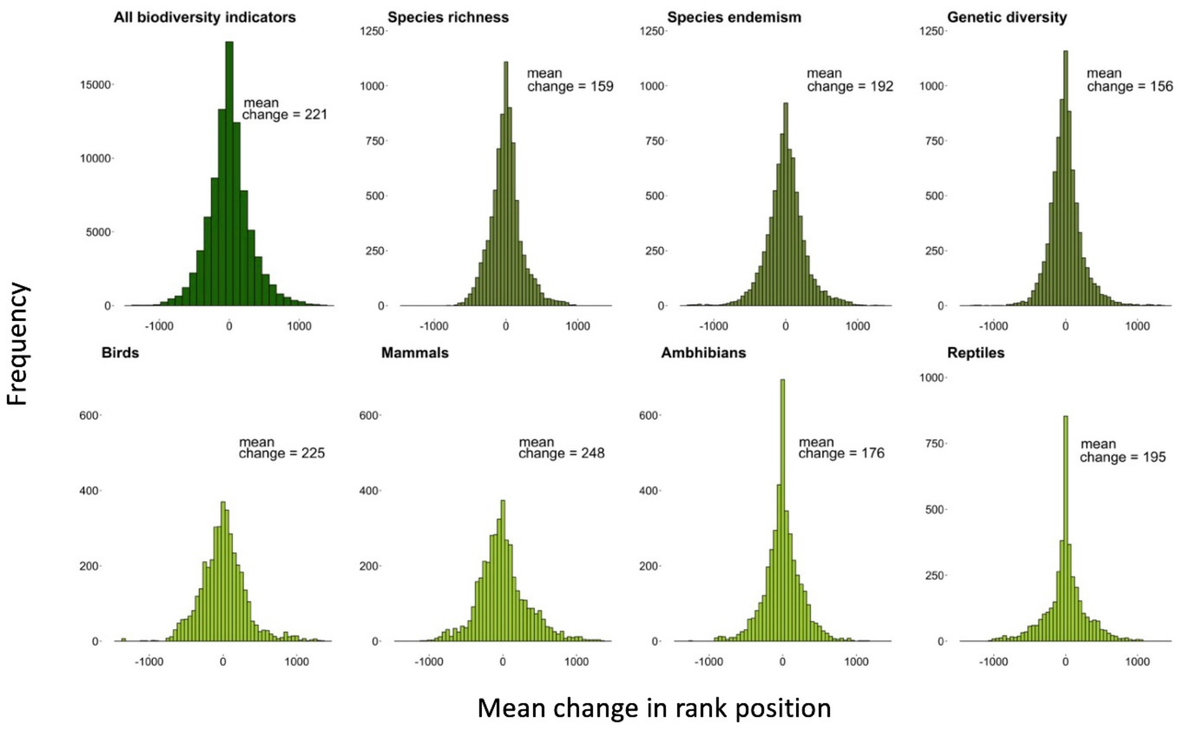

**Fig. S10:** Mean change in ranking position across all sites compared for all biodiversity indicators, for the three individual biodiversity indicators across all taxa and for the four taxa compared across all biodiversity indicators. To assess the mean change in rank position, all sites were ranked for each indicator and taxa individually and the average change in rank position per site was compared across the individual rankings (i.e. To assess the average change in rank position for species richness (SR) only, four rankings were compared: SR birds; SR mammals; SR; amphibians and SR reptiles. Subsequently the average change in rank position per site was calculated and plotted).

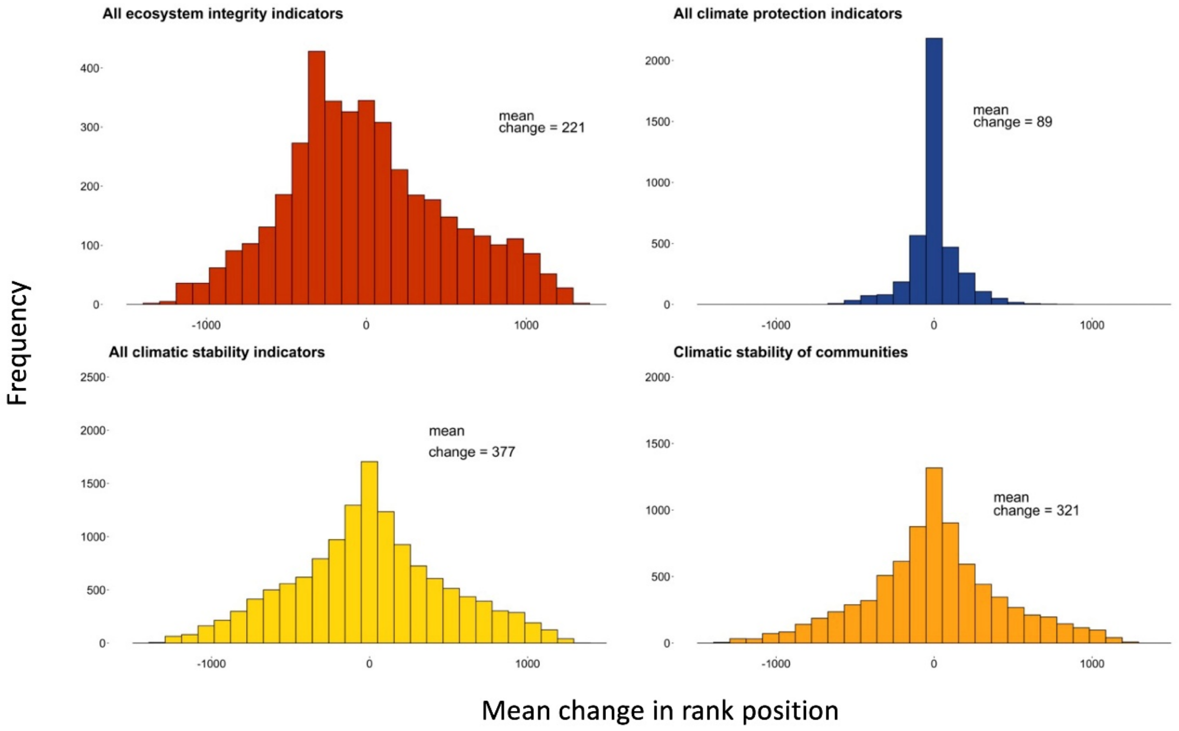

**Fig. S11:** Mean change in ranking position across all sites compared for all ecosystem integrity, climate protection and climatic stability. For climate stability the change in rank position across all indicators (climatic stability of species communities and change in forest cover) is shown in the bottom left graph and the change in rank position for climatic stability of species communities, considering the four included taxa individually, in the bottom right graph.

#### 2.6 The webinar

We introduced the site selection approach at a two-day online webinar, which was attended by 35 experts with a strong conservation background. During the workshop the different conservation objectives and indicator variables were presented and discussed. We used a questionnaire (Fig S12) to determine any missing conservation objectives or indicators as well as to allow everyone to order the conservation objectives by their perceived importance. In total 22 of the 35 attendants responded to the questionnaire.

**Conservation priority setting**

Please fill in the table below with a weighting of the different conservation strategies we introduced in the webinar session today. The weighting should be given from the perspective of your work sector. The weights should be allocated in the Legacy Landscapes context rather than based on other goals (e.g. regional or local development goals).

Weights allocated to the different conservation strategies should sum up to 100%. See example table in *Figure1*.

***By filling in this questionnaire, you agree that the data will be analyzed in anonymous form for a scientific publication.***

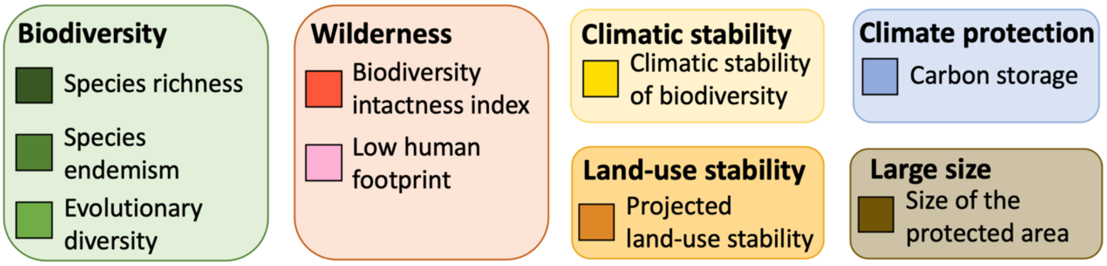

| Name | Biodiversity | Wilderness | Climatic stability | Land-use stability | Climate protection | Large size |
|---|---|---|---|---|---|---|
| … | **100%** | 0% | 0% | 0% | 0% | 0% |
| … | **50%** | **50%** | 0% | 0% | 0% | 0% |
| … | **50%** | 0% | **50%** | 0% | 0% | 0% |
| … | … | … | … | … | … | … |

High biodiversity

High wilderness

High climatic stability

*Figure 1: Example weighting table*

**Question 1.** Please fill in the weighting table from the perspective of your work sector, using percentages. Please use 5 percent intervals (e.g. 5%, 10%, 15%). If you filled in 'Other', please specify below the table.

| **Biodiversity** | **Wilderness** | **Climatic stability** | **Land-use stability** | **Climate protection** | **Large size** | ***Other*** |
|---|---|---|---|---|---|---|

If you filled in 'Other' please specify:

**Question 2.** Please (briefly) explain the motivation behind your weighting:

**Question 3.** Do we miss any important indicators within the six different conservation objectives (see Figure 1)? Please list:

**Question 4.** Do we miss any (macro ecological) conservation objectives in the Legacy Landscapes context (see Figure 1)? Please list:

**Question 5.** Which is the main work sector you would assign yourself to? Please choose one:

*Academia:*

*NGO:*

*Consultancy:*

*Government:*

*Other:*

*If other please specify:*

**Question 6.** Please identify your work place nationality (If you like to):

**Question 7.** Please identify your gender (m/f/d) (If you like to):

**Fig. S12:** The Questionnaire used during the workshop

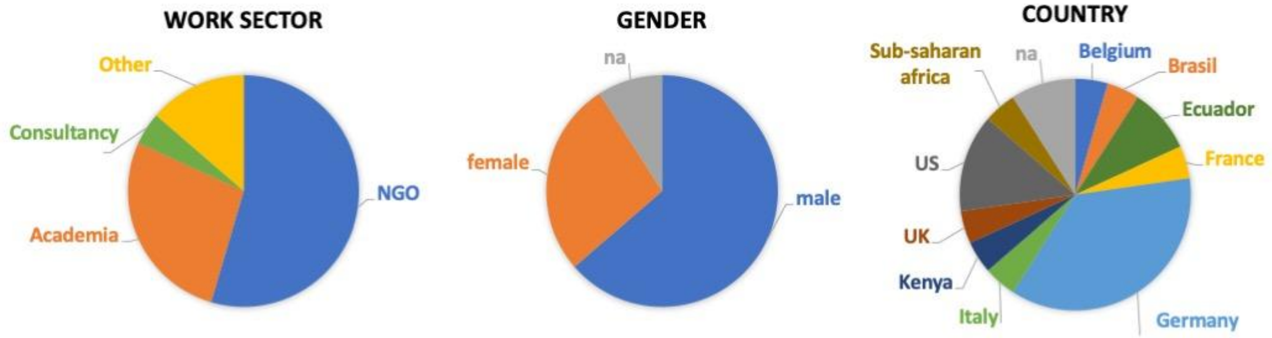

**Fig. S13:** Anonymous participant data for all workshop attendants who responded to the questionnaire.

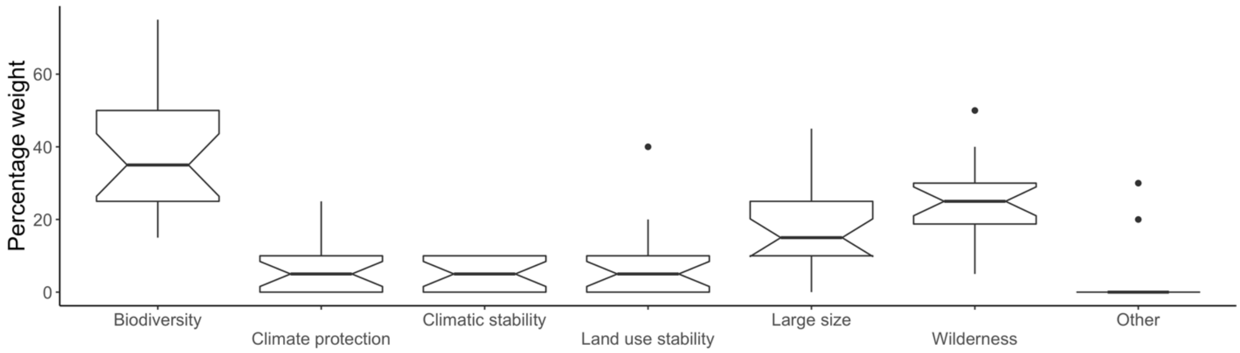

**Fig. S14:** Responses to Q1 by all attendants who filled in the questionnaire. Weights were allocated in 5 percent intervals to the individual objectives. Combined weights for all conservation objectives allocated per person summed up to 100 percent. Other included governance, ecosystem loss rate and socio-economic factors.

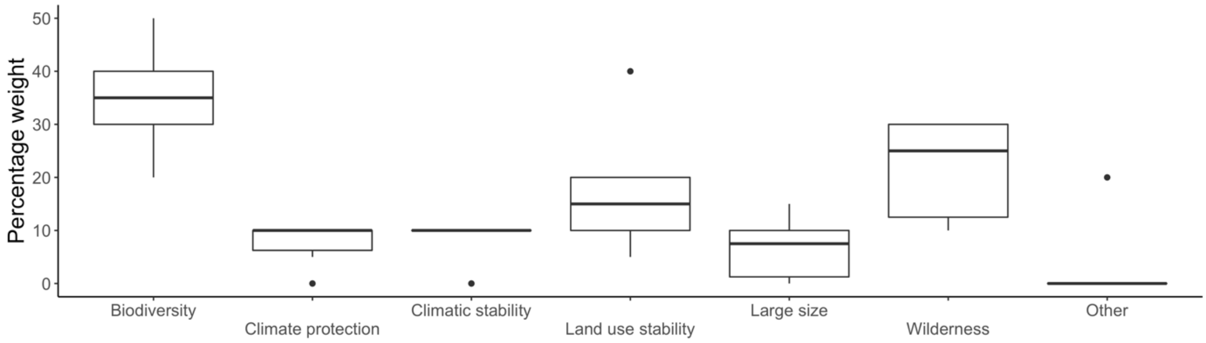

**Fig. S15:** Responses to Q1 by all attendants who described they work sector as academia. Weights were allocated in 5 percent intervals to the individual objectives. Combined weights for all conservation objectives allocated per person summed up to 100 percent (Other included socio-economic factors).

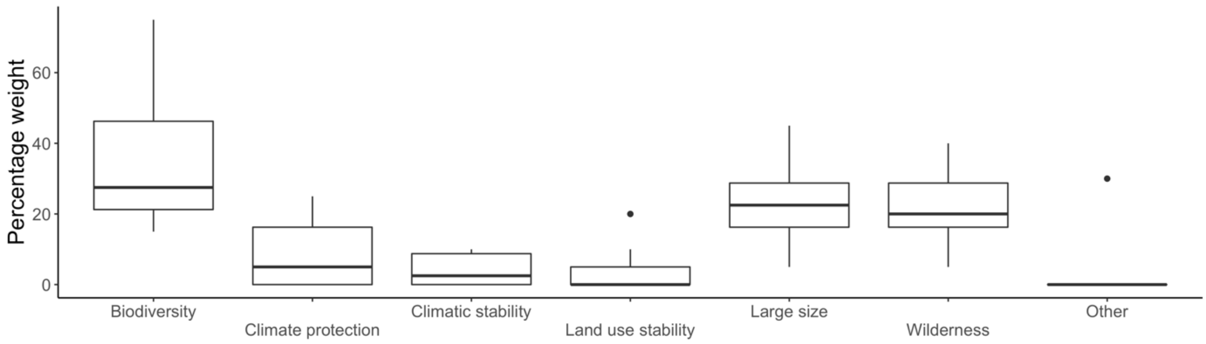

**Fig. S16:** Responses to Q1 by all attendants who described their work sector as NGO. Weights were allocated in 5 percent intervals to the individual objectives. Combined weights for all conservation objectives allocated per person summed up to 100 percent (Other included governance and ecosystem loss rate).

#### 3. Caveats

There are several limitations that need to be acknowledged when using the site selection tool with the current indicator. First, the biodiversity variables are calculated from global range maps of each terrestrial vertebrate species, which come at a coarse resolution, are of varying quality across species and taxa, and are therefore used for analysis at a 0.5° resolution; these cannot be used to derive accurate species lists for a given protected area [40]. Therefore, the included biodiversity variables give an indication of the biodiversity value of the region where a site is located, rather than accurate values for the individual site.

Second, there is always a high level of uncertainty surrounding any land-use and climate projections, which applies also to the models used to compute the indicators. Aside from specific, model-related uncertainties, the projected future impacts will largely depend on socioeconomic decisions and climate mitigation efforts [41]. Nevertheless, we believe that the large-scale geographic patterns of variables included in the analysis remain robust to these uncertainties and allow for a comparison across sites at the chosen resolution.

Next, to keep handling the decision support tool easy and thus allow a wider range of people to be able to use it, weights can only be applied to the individual conservation objectives. This results in limited possibilities to fine tune the evaluation of sites. Future versions of the tool will focus on adding more flexibility to the evaluation by adding additional options for more proficient users. These should include the possibility to weigh the individual indicators contained within the different conservation objectives.

Finally, the case study presented here is based on current biogeographic datasets. The tool developed allows for the preliminary evaluation of potential candidate sites for initiatives such as the LLF. Although the included datasets represent the state-of-the-art macro-scale data and allow for global as well as realm-wise comparisons across candidate sites, they cannot replace detailed on-the-ground evaluation of the individual sites.

### 4. The decision support tool

The decision support tool was developed to allow easy access to the different biogeographic datasets. It consists of four tabs and a settings panel on the left-hand sites which are described below:

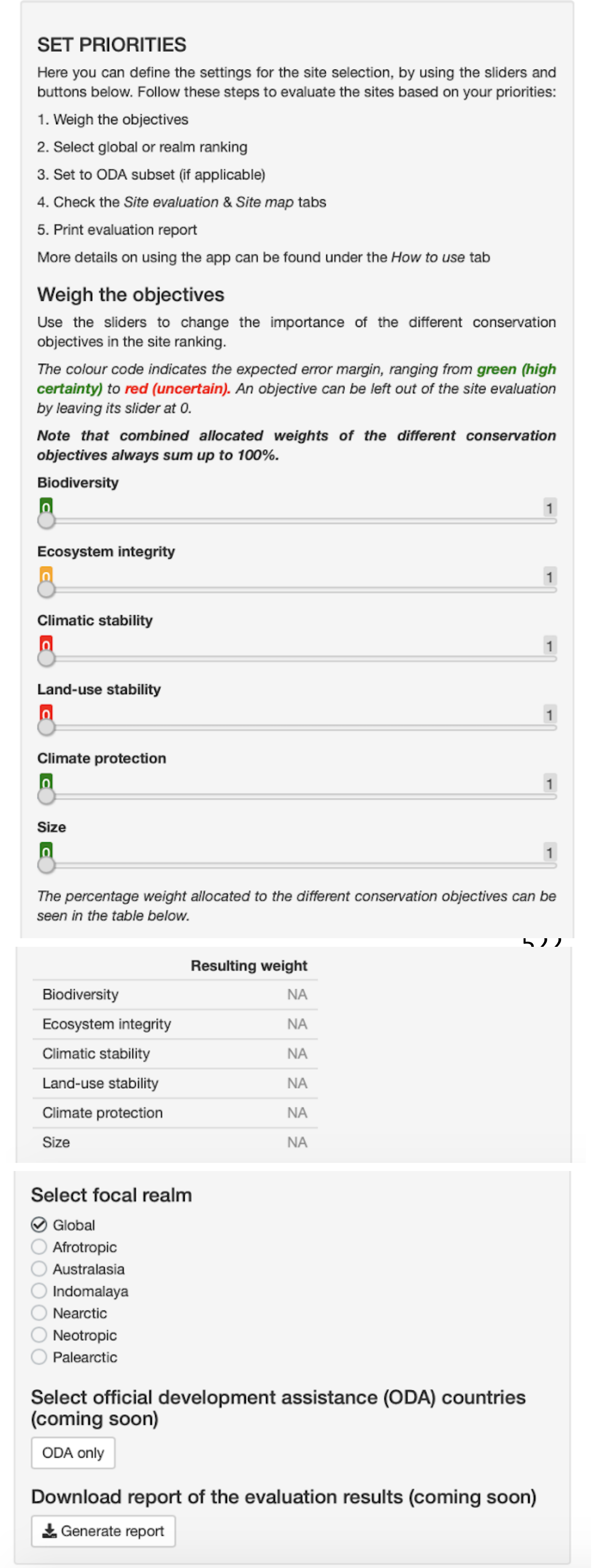

**Fig. S17:** The settings panel. The brief step by step instruction at the top gives a summary on how to use the conservation decision support tool. The sliders allow users to manually adjust the weighting of the individual conservation objectives (top).
The resulting allocated percentages can be seen in the tables below the sliders (center).
Below the weights table the user can select if sites should be selected globally or for a specific realm. With the “Select focal realm” button users can choose between evaluating sites globally or for one specific realm (bottom). The “Select official development assistance” button allows us to subset if all sites should be included in the evaluation or if only sites located in ODA countries should be included (bottom). The “Generate report” button allows downloading the generated evaluation based on the manually set weights and the selection of region and sites (bottom).

Background | Conservation objectives | Site evaluation | Site map | How to use

**The decision support tool**

The establishment and maintenance of protected areas (PAs) is viewed as a key action in delivering post 2020 biodiversity targets and reaching the sustainable development goals. PAs are expected to meet a wide range of objectives, ranging from biodiversity protection to ecosystem service provision and climate change mitigation. As available land and conservation funding are limited, optimizing resources by selecting the most beneficial PAs is therefore vital.

This decision support tool enables a flexible approach to PA selection on a global scale, which allows different conservation objectives to be weighted and prioritized. It is meant to facilitate a first evaluation of the potential of PAs for long term conservation before following up with detailed on the ground assessments of the candidate sites.

The current version of the decision support tool contains a set of 1347 PAs. The included PAs were selected as a case study subset based loosely on the criteria of the Legacy Landscapes Fund.

**Legacy Landscapes Fund**

Legacy Landscapes is a new international public-private initiative, led by the German Government, to develop and implement a conservation and financing strategy for the safeguarding of selected protected areas. The Legacy Landscapes Fund has a terrestrial focus and will significantly contribute to the post-2020 Biodiversity-Framework of the COP 15 at the CBD in 2021.

**The Legacy Landscapes Fund concept is based on three pillars:**

**1.** Permanent, stable and performance-based **funding** ensured by a combination of private donors and public funds of about one million $ per site per year.

**2.** Effective and efficient **management** that will be caried out in cooperation with national authorities, local communities and an NGO, with the annual disbursement of funds being controlled by an independent platform, and based on the fulfillment of certain indicators, the key performance indicators.

**3.** Strategic site selection, based on the **biogeography** of the site.

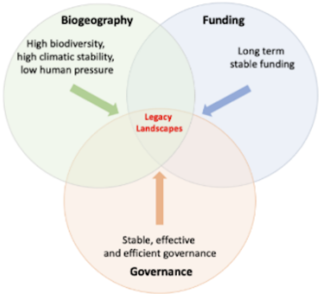

**Figure 1: The three cornerstones of the Legacy Landscapes Fund concept**

This decision support tool has been developed in cooperation between the Frankfurt Zoological Society and the Senckenberg Biodiversity and Climate Research Centre to support the selection of suitable sites for the Legacy Landscapes Fund. The tool enables the comparison of potential sites based on macro-ecological variables and thus falls under the Biogeography cornerstone of the Legacy Landscapes Fund concept. It facilitates the ranking of sites based on their performance across six different conservation objectives:

**Biodiversity, Ecosystem integrity, Climatic stability, Land-use stability, Climate protection and Size.**

Contact

**

FRANKFURT ZOOLOGICAL SOCIETY

Frankfurt Zoological Society

SENCKENBERG
world of biodiversity

Senckenberg Biodiversity and Climate Research Centre

**Fig. S18:** The “Background” tab of the conservation decision support tool. Here the user finds a brief introduction to the tool and its purpose.

Background | Conservation objectives | Site evaluation | Site map | How to use

**The conservation objectives**

Six conservation objectives were selected to enable the comparison of protected areas and evaluate their potential for long term conservation based on macroecological data. The objectives were chosen to allow a first assessment based on the size of the site, the biodiversity it contains, its intactness and its potential for future persistence. Each of the conservation objectives is measured based on one or more macro-ecological variables as described below:

**Biodiversity:** includes the richness, endemism and evolutionary diversity of species comprising four different taxa (mammals, birds, reptiles and amphibians)

**Ecosystem integrity:** includes the Biodiversity Intactness Index (BII), the human footprint and the observed change from biome to anthrome (change from natural to human modified landcover) in the area over the past 20 years

**Climatic stability:** includes the projected impacts of climate change on the stability of ecological communities (mammals, birds, reptiles and amphibians) and the change in tree cover by the mid of the century under a medium warming scenario

**Land-use stability:** includes the projected change in land-use in the buffer zone around the site

**Climate protection:** includes the amount of baseline, vulnerable and irrecoverable carbon storage within the site, indicating the contribution of the site to climate protection by binding carbon dioxide.

**Size of the site:** is the extent of the site in km2

The six different conservation objectives can be combined into different conservation goals as laid out in the figure below. Using the sliders on the left the conservation objectives can be combined by allocating a weight to each objective. Objectives allocated a weight of 0 are excluded from the weighting. The resulting ranking based on the selected objectives and the allocated importance (weight) can be seen in the ***Site evaluation*** tab. The location of the top scoring sites can be seen in the ***Site Map*** tab.

*Details on the included variables and data sources and can be found in the text box at the bottom of the page.*

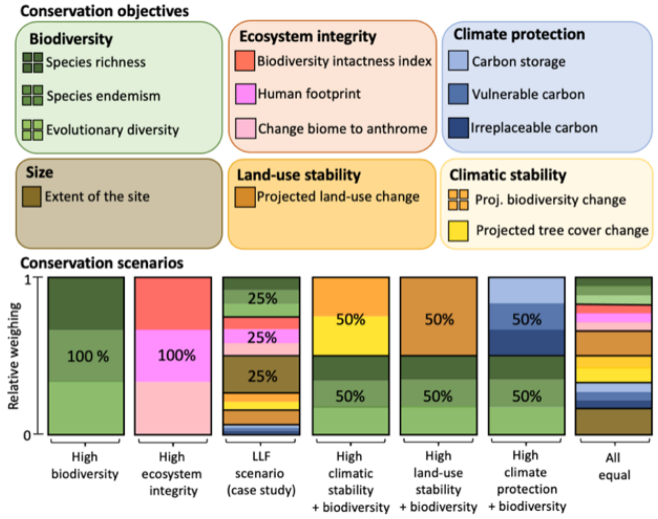

**Figure 2: Conservation objectives and strategies** The six different conservation objectives Biodiversity, Ecosystem integrity, Climatic stability, Land-use stability, Climate protection and Size can be combined into different conservation scenarioes. These conservation scenarioes allow to weigh the different conservation objectives against each other, to set priorities when evaluating sites for conservation.

**Fig. S19:** The "Conservation objectives" tab gives the user an overview over the six conservation objectives included in the conservation decision support tool and the indicators they consist of. At the bottom of the tab the user can find a PDF that explains the included data in greater detail (the content of the PDF can be found below under *3.1.1 Details on the conservation objectives*).

Background | Conservation objectives | Site evaluation | Site map | How to use

Site evaluation based on weighted objectives

The ranking table shows the overall ranking of the potential sites based on the applied weights. The values for the different conservation objectives are scaled across all sites included in the ranking. The values range from 0 to 1, with 0 being allocated to the site with the lowest score and 1 being allocated to the site with the highest score for the conservation objective. The scaled ranks are shown for each conservation objective, for each site on the right hand site of the table and remain the same independent of the weighting. This means the scaled value that a site has for a certain conservation objective indicates the overall ranking position of that site for that objective, as the following example shows:

**If the weights for 'Biodiversity' and 'Ecosystem integrity' are both set to 50%, you will see that the 'Talamanca Range' is the top site. This is because it has the second highest biodiversity among the included sites, with a 'Biodiversity' score of 0.99. But it also has a clear human footprint, indicated by the 'Ecosystem integrity' score of 0.71. In comparison the combined site 'Manu - Alto Purus' ranks third globally with a very good 'Biodiversity' score of 0.70 but additionally it is also very pristine with a very high 'Ecosystem integrity' score of 0.93.**

*The ranking table can be adjusted by using the sliders on the left hand side. Allocating different weights to the individual objectives will change the ranking of the sites. Using the* ***Select focal realm*** *buttons above the table, the ranking table can be subset to show the resulting ranking for the individual realms or across all sites globally.*

**Show** 10 **entries** **Search:**

| | Rank | Realm | International Name | Biodiversity | Ecosystem integrity | Climatic stability | Land-use stability | Climate protection | Size |
|---|---|---|---|---|---|---|---|---|---|
| 1 | 1 | Australasia | Lorentz | 0.607396086 | 0.80230815 | 0.548465231 | 0.654322973 | 0.834105246 | 0.073520033 |
| 2 | 2 | Australasia | Telefomin | 0.607252524 | 0.912244933 | 0.537948473 | 0.685210529 | 0.810333302 | 0.006225128 |
| 3 | 3 | Australasia | Wet Tropics of Queensland | 0.520467601 | 0.741410564 | 0.807227266 | 0.889916118 | 0.263765134 | 0.027954551 |
| 4 | 4 | Australasia | Enarotali | 0.507679885 | 0.819943063 | 0.576880068 | 0.658567367 | 0.796953488 | 0.008785558 |
| 5 | 5 | Australasia | Pegunungan Wayland | 0.446090326 | 0.865420475 | 0.61295482 | 0.658441202 | 0.862794401 | 0.00430076 |
| 6 | 6 | Australasia | Pegunungan Tamrau Selatan | 0.419463833 | 0.885203088 | 0.784841706 | 0.625291729 | 0.860900299 | 0.014886721 |
| 7 | 7 | Australasia | Girringun | 0.407186504 | 0.67611454 | 0.858131928 | 0.896210488 | 0.26227229 | 0.008657429 |
| 8 | 8 | Australasia | Pegunungan Tokalekaju | 0.365977228 | 0.776801303 | 0.571428877 | 0.593722704 | 0.569596224 | 0.012244087 |
| 9 | 9 | Australasia | Gondwana Rainforests of Australia | 0.345034756 | 0.689581806 | 0.834040409 | 0.803964452 | 0.373084456 | 0.011536259 |
| 10 | 10 | Australasia | Bogani Nani Wartabone | 0.324413688 | 0.676782772 | 0.674263254 | 0.642144486 | 0.572940405 | 0.008879256 |

Showing 1 to 10 of 1,345 entries — Previous 1 2 3 4 5 … 135 Next

**Fig. S20:** The "Site evaluation" tab shows the evaluation results based on the set weights and selected region and type of sites (ODA or not) in a table. Sites are ranked from performing best to least under the respective settings.

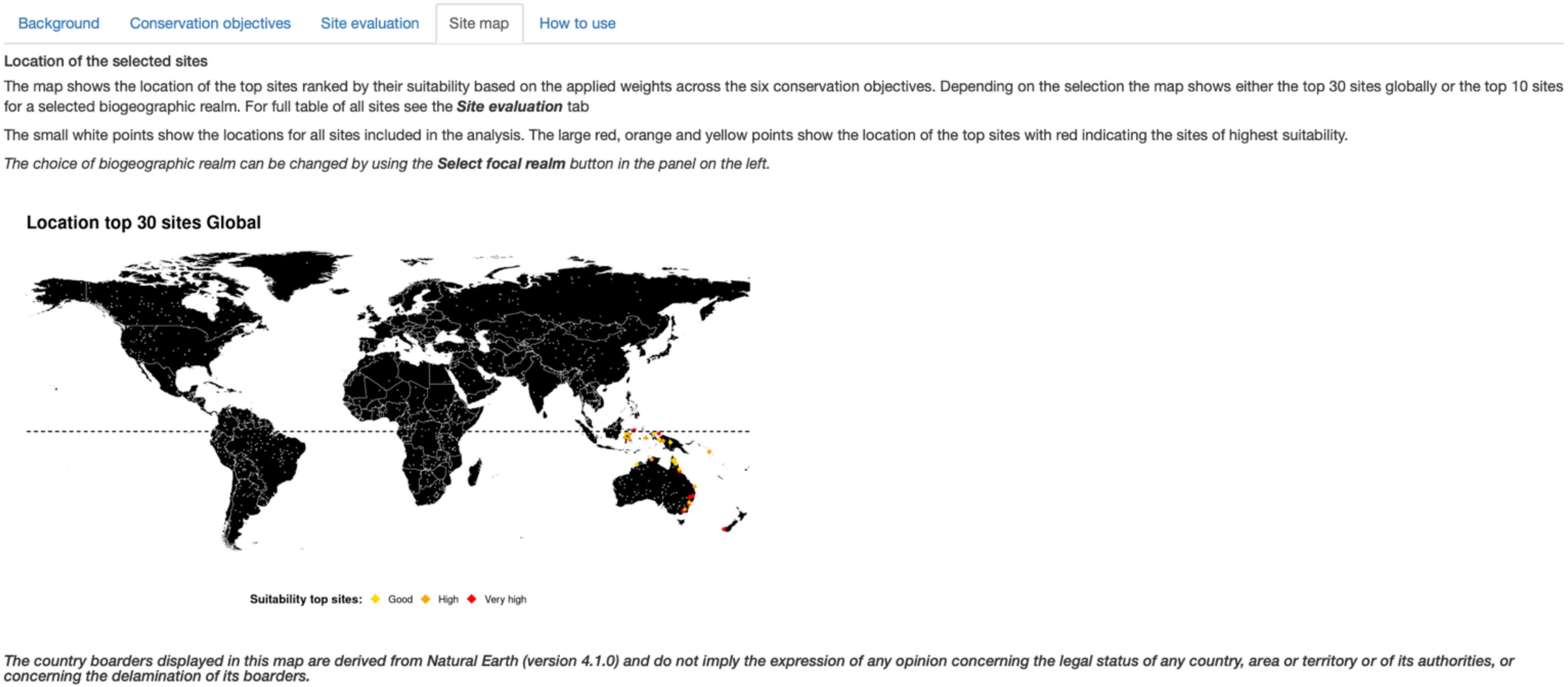

**Fig. S21:** The “Site map” tab shows the spatial distribution of the top 30 sites based on the set weights and selected region and type of sites (ODA or not).

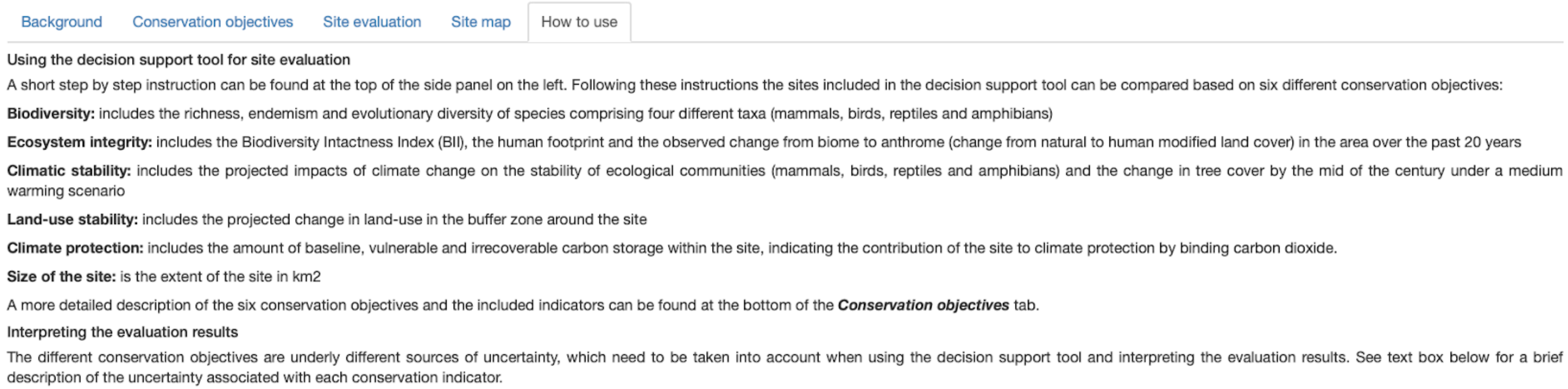

**Fig. S22:** At the bottom of the tab the user can find a PDF with more detailed instructions and information on how to interpret the results and the uncertainty around the different objectives (the content of the PDF can be found below under *3.1.2 How to use the conservation decision support tool*).

#### 4.1 User manual decision support tool

To help users understand the datasets underlying the decision support tool and enable them to use the tool to evaluate sites for conservation, the tool includes a brief description of the included data and a user manual.

##### *4.1.1 Details on the conservation objectives*

*The site data*

The sites currently included in the conservation decision support tool are all registered sites under either one or more of the following criteria:

- a protected area from the global world database in protected areas [1] that is listed by the International Union for Conservation of Nature (IUCN) in either category I or II,
- a natural World Heritage Site (WHS),
- a Key Biodiversity Area (KBA).

The shapefiles for the IUCN protected areas as well as the World Heritage Sites were derived from protected planet [1]. The Shapefiles for the KBAs were derived from BirdLife International [2].

*The conservation objectives data*

The six different conservation objectives which are included in the decision support tool are biodiversity, ecosystem integrity, climatic stability, land-use stability, carbon storage and size. Each of these objectives consists of one or several underlying macro-ecological indicator variables. See below for a detailed description of the variables included within each of the six conservation objectives and how these variables are derived (*Shorter and simpler explanations can be found under the tab "**How to use**"*).

*Biodiversity*

The biodiversity objective includes three different variables, the total number of species, the degree of endemism and the evolutionary diversity of the species occurring in the region the site is located in.

Species richness

The species richness, for four taxa of vertebrates, is derived from range maps for virtually all species of the four terrestrial vertebrate taxa: from the BirdLife International for birds [4], the IUCN for mammals and amphibians [3], and from GARD for reptiles [5].

*Sites with a higher species richness are allocated a higher suitability for long-term conservation than sites with a lower species richness.*

*Endemism*

To capture biodiversity that is unique to a region, a measure for the prevalence of range restricted (endemic) species within the region is used. Species endemism is estimated by calculating weighted range size rarity, which is the sum of the inverted range extents of all species, divided by the number of species occurring in a site [6].

*Sites with a higher rate of species endemism are allocated a higher suitability for long-term conservation than sites with a lower rate of species endemism.*

*Evolutionary diversity*

Evolutionary diversity is included to have an estimate of how evolutionary unique the species within a region are. Measures of evolutionary diversity can give an idea of how much evolutionary history is stored within a set of species. A high amount of evolutionary history might imply a high feature diversity across the species within the region and could, arguably, make a community more resilient to disturbance. Evolutionary diversity is calculated using phylogenetic endemism (PE), which is a combined measure of evolutionary history and the uniqueness of a species community. PE identifies regions with high numbers of evolutionary isolated and geographically restricted species. In addition to summing the shared evolutionary history of a species assemblage, PE also incorporates the spatial restriction of phylogenetic branches covered by the assemblage [12].

*Sites with a higher evolutionary diversity are allocated a higher suitability for long-term conservation than sites with a lower evolutionary diversity.*

*Ecosystem Integrity*

The ecosystem integrity objective includes three different variables, the biodiversity intactness index (BII), the human footprint in and around the site and the change from biome to anthrome in the past two decades.

*Biodiversity intactness index (BII)*

The BII presents the modeled average abundance of present species, relative to the abundance of these species in an intact ecosystem [16]. This means the index gives an indication of how much species abundances in a region have already changed due to anthropogenic impacts e.g. land-use change. For the BII we are using the global map of the Biodiversity Intactness Index calculated by Newbold et al (2016).

*Sites with a higher estimated biodiversity intactness are allocated a higher suitability for long-term conservation than sites with a lower biodiversity intactness.*

*Human footprint*

As a measure of how pristine the sites still are, a measure of the human footprint within the region is included. Estimates of the human footprint within sites are derived from the standardised human footprint layer by Venter et al (2016), which includes data on the extent of built environments, crop land, pasture land, human population density, night-time lights, railways, roads and navigable waterways.

*Sites with a lower human footprint are allocated a higher suitability for long-term conservation than sites with a higher human footprint.*

<u>*Land-use change*</u>

To derive past changes in the land cover of a site we calculated the average percentage change across the site from biomes (natural vegetation cover) to anthromes (human-modified land cover such as rainfed cropland, irrigated cropland, mosaic cropland, mosaic natural vegetation and urban areas). The fraction of land cover classes time series, ranging from 1992 – 2018, was obtained from the GEOEssential project [18].

*Sites with a lower percentage of land-use change are allocated a higher suitability for long-term conservation than sites with a higher percentage of land-use change.*

*Climatic stability*

The climatic stability objective consists of two different variables: the projected stability of animal biodiversity and the projected tree cover change under future climate change.

<u>*Climatic stability of biodiversity*</u>

To estimate the climatic stability of a site we are looking at the potential impacts of climate change on the biodiversity within the site. Climate change is driving shifts in species distributions and it is well established that many taxa are shifting their ranges towards higher latitudes and elevations. But also, idiosyncratic species responses to climate change have been observed. These heterogeneous range shifts have the potential to reshuffle species assemblages, which can have highly unpredictable impacts on species interactions and ecosystem functions (e.g., changes in prey predator relationships or competition). We assume that species assemblages that are not predicted to change a lot in future or experience large species losses are under less risk from climate change than species assemblages that experience a lot of reshuffling. Therefore, we include projected turnover in species under future climate change as an indicator for the climatic stability of biodiversity. Projections of species ranges are derived from species distribution models (see Hof et al 2018 for a detailed description of the modelling). For each site all species that are projected to occur there currently and/or in future (2050) are extracted. The turnover is then calculated between the current and future species assemblage of a site, using the formula for Bray Curtis dissimilarity [28].

*Sites with higher climatic stability (i.e., a lower projected turnover in species) are allocated a higher suitability for long-term conservation than sites with a lower climatic stability.*

*Forest cover change*

We included the projected change in tree cover derived from the LPJ-GUESS process-based dynamic vegetation-terrestrial ecosystem model [29]. The climate input for the model was derived from the ISIMIP2b simulations, described above under climatic stability of biodiversity. The projected change of tree cover is calculated as the average percentage change projected to occur within the site.

*Sites with a lower change in the projected tree cover are allocated a higher suitability for long-term conservation than sites with a higher change in projected tree cover.*

*Land-use stability*

To assess the potential impacts of projected future land-use change we used predictions of the change in pastures, croplands and biofuel croplands in the buffer zone around the sites (50 km buffer), excluding the site itself.

*Projected land-use change*

Projected land–use change is derived from simulations of current and future land-use, based on global land-use change models, using the assumptions of population growth and economic development as provided by ISIMIP2b and described in Frieler et al. (2017). The used land-use change models [30,32] account for climate impacts (e.g., on crop yields) and were driven with the same climate input as the species distribution models used to derive climatic stability of biodiversity (see above). The land-use scenarios provide percentage cover of six different land-use types (urban areas, rainfed crop, irrigated crop, pastures, as well as rainfed and irrigated bioenergy crops). We averaged annual land-use data for each of two different time periods (1995 and 2050), across the four GCMs (see above under Climatic stability), and calculated a combined value of average land-use change for the buffer zone around each site.

*Sites with a lower projected increase in land-use in the buffer zone are allocated a higher suitability for long-term conservation than sites with a higher projected increase in land-use in the buffer zone.*

*Carbon storage*

The carbon storage objective includes three different variables, using the three dimensions of ecosystem carbon stocks as defined by Goldstein et al. (2020). These include the amount of manageable carbon stocks that currently exist but could be influenced in principle by human actions, the amount of vulnerable carbon stocks that currently exist and will be released if land-use changes and the amount of irrecoverable carbon stocks in a site.

*Manageable carbon*

As an indicator for the climate protection capacity, we used the estimated amount of manageable carbon as provided by Noon et al (2021). This layer includes the amount of carbon stored in the above and below ground vegetation as well as soil organic carbon stocks up to 30 cm depth, or up to 100 cm within inundated soil, as these depths are most relevant to common disturbances [34]. We derived the average amount of carbon in t per ha for each site.

*Sites with higher baseline carbon stocks are allocated a higher suitability for long-term conservation than sites with lower baseline carbon stocks.*

*Vulnerable carbon*

Vulnerable carbon is defined by Goldstein et al (2020) as the amount of the manageable carbon, described above, that is likely to be released through typical land conversion in an ecosystem. We derived the average amount of vulnerable carbon in t per ha for each site.

*Sites with higher vulnerable carbon stocks are allocated a higher suitability for long-term conservation than sites with lower vulnerable carbon stocks.*

*Irrecoverable carbon*

Irrecoverable carbon is defined as the amount of the vulnerable carbon, described above, that if it is lost through typical land conversion actions, cannot be recovered over the following 30 years [34]. We derived the average amount of irrecoverable carbon in t per ha for each site.

*Sites with higher irrecoverable carbon stocks are allocated a higher suitability for long-term conservation than sites with lower irrecoverable carbon stocks.*

*Large size*

For the extent of the area, we preselected sites that are larger than 2000 $km^2$, based on the precondition that Legacy Landscapes should have a minimum size to maintain a viable ecosystem.

*Extent of the site*

The area in $km^2$ is derived from the site polygons provided by protected planet [1] or the Key Biodiversity Area (KBA) database [2].

*Larger sites are allocated a higher suitability for long-term conservation than smaller sites.*

*4.1.2 How to use the conservation decision support tool*

The conservation decision support tool is meant to facilitate global or realm wise comparisons of sites based on macroecological datasets. The spatial scale of the included datasets enables the user to compare a vast number of sites globally based on the six different conservation objectives. Nevertheless, two important points need to be kept in mind when using the decision support tool and interpreting the evaluation results.

*Large-scale comparison, not local assessment*

Firstly, due to the coarse resolution of most globally available datasets the decision support tool facilitates a first evaluation of the included sites but *should not be used for local assessments*. This means that for the selection of specific areas for conservation and the practical implementation of nature conservation on the ground requires further evaluation steps that a tool like this cannot cover. These further steps should involve an on-site assessment based on additional parameters at a higher resolution (e.g. more detailed biological data acquired through surveys and observations). For a final decision, it is also crucial to consider non-biological characteristics, ranging from available infrastructure, NGO presence, political situation, access to the site and potential funding possibilities to socio-economic factors.

*Underlying data uncertainty varies among objectives*

Secondly, the different indicator datasets included within the six conservation objectives come with different levels of uncertainty and error margins, which affects the resulting ranking. These varying error margins should be kept in mind when interpreting the results. For example, a ranking of sites based exclusively on the biodiversity objective is less prone to errors, because the global patterns of species richness and diversity are well-known and unlikely to change substantially in the near future at the used spatial scale. In contrast, the climatic stability objective is based on modelling of future biodiversity responses to climate change, which are sensitive to human societal and political decisions and need to be regularly updated with ongoing developments and new knowledge; therefore, the ranking of sites based exclusively on the climatic stability objective is more prone to errors and could change in the future. *We have therefore colour-coded the sliders for the individual objectives in the panel on the left based on the expected error margin, ranging from* ***green*** *(high certainty) via* ***yellow*** *(intermediate certainty) to* ***red*** *(uncertain). An objective can be left out entirely of the site evaluation by leaving its slider at 0.* Below we briefly describe the underlying main sources of uncertainty that should be considered with each conservation objective.

*Biodiversity objective:* Low error margin

This objective consists of three conservation indicators:

- species richness is the number of species occurring in the region the site is located in and is derived from species range polygons provided by BirdLife International (birds [4]), IUCN (mammals, amphibians [3]) or GARD (reptiles [5]).

- endemism is the range size rarity across all species occurring within the site.

- evolutionary diversity is calculated using phylogenetic endemism (PE), which is a combined measure of evolutionary history and the uniqueness of a species community. PE identifies areas with high numbers of evolutionary isolated and geographically restricted species.

The base data for these indicators are globally available species range maps for virtually all species in the four classes of terrestrial vertebrates (mammals, birds, reptiles, and amphibians) and, for evolutionary diversity, phylogenies that describe how species are related to each other. The observed indicator patterns are well-known and therefore stable at the global scale and unlikely to introduce high amounts of uncertainty into the site evaluation, although we acknowledge that the individual species range maps are only rough representations of where species actually occur and should therefore not be used for local assessments. Similarly, some uncertainty exists in the phylogenetic tree. Due to the coarse nature of the range maps, the resulting species numbers for the individual sites should be interpreted as the number of species occurring within the region where the site is located, not as the exact number of species known to occur within the site.

*Ecosystem integrity objective: Intermediate error margin*

The ecosystem integrity objective includes three conservation indicators with differing error margins:

- The biodiversity intactness index (BII) connects modelled land-use pressures on biodiversity with locally observed biodiversity data from the PREDICTS project. There are several sources of uncertainty associated with this modelling approach, including the quality of the underlying biodiversity data and the modelling approach itself. We therefore consider the error margin for this conservation indicator as higher compared to e.g. the indicators included in the biodiversity or size objective, but not as high as the completely modelled indicators such as climatic stability. Details on the BII can be found in Newbold et al 2016.

- The human footprint (HFP) within the sites was estimated using the data of Venter et al (2016). The standardized HFP provided by the source data includes the extent of built environments, cropland, pasture land, human population density, night-time lights, railways, roads and navigable waterways. Data included in the footprint dates partially back to 2009 and might not reflect recent developments within and around the actual sites. Therefore, we consider the error margin for this indicator to be higher compared to e.g. the indicators included in the biodiversity or size objective, but not as high as the completely modelled indicators such as climatic stability.

- The biome to anthrome change over the last 20 years measures the conversion of natural ecosystems to different human-dominated land-use categories. This indicator is derived from satellite pictures, which are classified into biome and anthrome classes [18]. From these classes, the percentage change in class coverage across the image pixels falling into each site is then calculated. This indicator has a low error margin, as it is unlikely to introduce high amounts of uncertainty into the site evaluation.

*Climatic stability objective: High error margin*

The climatic stability objective includes two conservation indicators with high error margins:

- projected change in biodiversity until 2050 modelled under a medium emission pathway (IPCC scenario RCP 6.0 [42]) and associated level of global warming
- projected change in tree cover until 2050 modelled under a medium emission pathway (IPCC scenario RCP 6.0 [42]) and associated level of global warming

Both indicators are based on models, which come with various sources of uncertainty, including the underlying biodiversity data, the chosen model type and the climatic drivers and associated models (details on can be found here [24,29]). Projected change in biodiversity is the turnover in species community compositions between today and 2050 based on species-specific distribution models for virtually all species of the four classes of terrestrial vertebrates (mammals, birds, reptiles, and amphibians) projected onto modelled future climatic conditions. Projected change in tree cover is measured as the percentage change between today and 2050 based on a global dynamic vegetation model that was run for modelled present and future climatic conditions. These projections give an estimate where the impacts of climate change are expected to be severe and which areas might be less affected, but they come with high levels of uncertainty and models are constantly updated as they are based on human societal behaviour and political decisions. We thus expect a relatively high error margin for the climatic stability objective compared to the other objectives.

*Land-use stability objective: High error margin*

The land-use stability objective is based on one conservation indicator:

- percentage of projected land-use change in a buffer zone around each site (50 km buffer from site margin) until 2050 modelled under a medium emission pathway (IPCC scenario RCP 6.0 [42]) and associated level of land-use conversion [e.g. from pasture to cropland].

The underlying modelled data are matching those for the conservation indicators included in the climatic stability objective. These models come with several sources of uncertainty and additionally depend on the applied assumptions of population growth and economic development (details on the methods and potential sources of uncertainty can be found here [30,32]). The projected changes in land-use give an indication where circumstances might be beneficial for a future increase in land-use potentially adding additional pressures on sites, but these projections are highly uncertain and need to be constantly updated as they are based on human societal behaviour and political decisions. The expected error margin for the land-use stability is thus expected to be high.

*Carbon storage objective: Low error margin*

The carbon storage objective consists of three different measures of carbon storage as a conservation indicator:

- baseline carbon, i.e. the amount of carbon stored in the above and below ground as well as the soil organic carbon of an ecosystem.
- vulnerable carbon is defined as the amount of (baseline) carbon that is likely to be released through typical land conversion in an ecosystem.
- irrecoverable carbon, is defined as the amount of carbon, that if it is lost through typical land conversion actions, and that cannot be recovered over the following 30 years.

All three measures are derived from the same data source [33] and measure carbon storage because this effectively removes the greenhouse gas carbon dioxide (CO2) from the atmosphere, thus protecting the current climate system from global warming effects. The baseline carbon estimates for the underlying dataset have been derived from various sources and combine the best estimates available. Whilst the amount of vulnerable and irrecoverable carbon strongly depend on the estimates of carbon lost through land conversion and recovery time, the overall spatial patterns of carbon storage are well-known and likely to be stable. The expected error margin for the carbon storage objective is thus expected to be comparatively low, contrary to the climatic and land-use stability objectives which depend on complex modelled datasets.

*Size objective: Low error margin.*

The only conservation indicator for the size objective is the size of the sites. This is directly calculated from shapefiles provided by the World Database on Protected Areas [1] and BirdLife International [2] and has an expected low error margin. As the calculated size depends on the accuracy of the shapefiles, this accuracy might therefore slightly affect the site evaluation for some included sites, but the errors are likely to be minor.
